# Feasibility of PIANO-Cog for older adults: A randomised controlled pilot trial exploring changes in cognition and brain microstructure

**DOI:** 10.64898/2026.01.28.702269

**Authors:** Fionnuala Rogers, Carolyn McNabb, Ege Erdem, Claudia Metzler-Baddeley

**Affiliations:** Cardiff University Brain Research Imaging Centre (CUBRIC), School of Psychology, Maindy Road, Cardiff University, Cardiff, United Kingdom; Department of Engineering, Faculty of Natural, Mathematical & Engineering Sciences, King’s College London

**Keywords:** Ageing, cognitive intervention, piano training, neurologic music therapy, therapeutic instrument music performance, executive function, fluid intelligence, neuroplasticity, microstructure, MRI, pilot, feasibility, verbal fluency

## Abstract

**Background:** Executive functions are a key target of cognitive interventions for older adults due to their central role in daily functioning and maintaining a good quality of life. Piano training has been proposed as an ecologically valid method of improving cognition and brain structure in older adults. The primary aims of this study were to (i) evaluate the feasibility and acceptability of *Piano Instruction for Adult Novices as an Online Cognitive Intervention* (PIANO-Cog), a novel bespoke 8-week self-guided piano training programme for adults over 50 years of age, and (ii) assess the feasibility of conducting a fully-powered randomised controlled trial (RCT), including recruitment, retention, and adherence. Secondary aims explored effects of PIANO-Cog on executive functions and brain microstructure using advanced diffusion-weighted imaging (DWI).

**Method:** Thirty-three healthy music novices aged 51–80 years (M = 63.73, SD = 7.94) participated in a two-arm unblinded feasibility RCT. Participants were assigned by stratified allocation for age and sex to either (i) 8 weeks of PIANO-Cog, requiring 30 minutes of practice, 5 days per week, or (ii) a passive control group. Cognitive assessment and MRI scanning were conducted before and after the intervention using a strong-gradient (300mT/m) 3T Connectom scanner to acquire multi-shell DWI data with b-values ranging from 200 to 6,000 s/mm². Grey and white matter microstructure were modelled with Soma And Neurite Density Imaging (SANDI) and Neurite Orientation Density and Dispersion Imaging (NODDI).

**Results:** According to predefined criteria, feasibility was established for recruitment (91.6%), retention (75%) and adherence (>100%) rates. Preliminary observations suggest that piano training compared with control was associated with improvements in verbal fluency and multiple changes in brain microstructure including increases in apparent soma size and radius and reductions in extracellular signal in frontal and temporal cortical regions, larger apparent neurite density in right inferior frontal gyrus and changes in neurite dispersion in left middle temporal and right precentral gyri.

**Discussion:** The results demonstrate that short-term remote piano training is a feasible cognitive intervention for healthy adults over 50. Preliminary evidence suggest that PIANO-Cog was associated cognitive improvements and changes in brain microstructure in executive, auditory and motor regions.

## 1. Introduction

Normal cognitive ageing is associated with declines in fluid intelligence and executive functions (EF) (1), which are supported by prefrontal and fronto-striatal-cerebellar networks (2). These declines can impact independence and quality of life (3), making EF a key target for cognitive training interventions for healthy ageing. Computerised cognitive training programmes often fail to produce generalised (“far transfer”) effects on everyday functioning (4–7), likely due to their limited ecological validity (8). EF are hypothesized to have developed as an extension of the motor control system for optimal interaction with the environment through predictions of stimulus-response associations regulated by the basal ganglia and movement control through predictive-error feedback loops by the cerebellum (9). Therefore, activities which engage EF through motor and sensory integration are now recommended for cognitive interventions (10).

Music-based activities, particularly instrument learning which involves generating and anticipating the outcomes of movements, engage widespread sensorimotor, auditory, and executive networks (11), providing a more ecologically valid form of cognitive stimulation. The basal ganglia and cerebellum have distinct roles in both the interpretation of rhythm and EF (12–14). Older adults who play a musical instrument were reported to have a reduced risk of developing dementia in cross-sectional, longitudinal and twin studies (15–17). In a systematic review and meta-analysis of 502 participants across 13 studies, we found a moderate effect on processing speed (*d =* .47, *p <* .0001), a low-moderate effect on attention-switching (*d =* .39, *p =* .002) and inhibitory control (*d =* .39, *p =* .034) in older non-musicians who learned a new instrument compared to an active or passive control group (18).

Neuroanatomical differences have been consistently observed between musicians and non-musicians in brain regions involved in EF, auditory and motor processing. Specifically, voxel-based morphometry (VBM) and cortical thickness (CT) studies have shown greater grey matter volume or thickness in musicians in Heschl’s gyrus, the planum temporale, fusiform gyrus, cerebellum, superior parietal lobule, inferior frontal lobe, lingual gyrus, hippocampus, and caudate nucleus (19–25). White matter pathways connecting these regions also show microstructural differences in musicians. For example, musicians exhibit greater fractional anisotropy (FA; a measure of diffusion coherence used as a proxy for white matter integrity) (26), in the arcuate fasciculus (27) – a key white matter tract linking temporal and frontal regions, important for the integration of motor actions with sounds (28), - as well as in the anterior corpus callosum and corticospinal tract (29), which are integral for bimanual coordination and motor skill execution. In line with cross-sectional findings, a recent trial, *Train the Brain with Music* (30), has demonstrated structural changes in Heschl’s gyrus and caudate nucleus, and maintenance of the fornix microstructure (31) following music training in healthy older non-musicians using CT, VBM and fixel-based analysis measurements, respectively (32,33).

Taken together, there is accumulating evidence that musical training, such as learning to play the piano, has potential for combating cognitive ageing by helping to maintain EF and fluid abilities and by slowing age-related neural decline. Based on this evidence, we developed a novel bespoke self-guided piano training programme, Piano Instruction for Adult Novices as Online Cognitive Intervention (PIANO-Cog), that allows remote practice at home. Details of the PIANO-Cog intervention and the protocol of the present feasibility RCT have previously been published (34). In short, the PIANO-Cog program is an 8-week home-based piano training intervention for older adults, comprising eight 20-minute instructional videos and an accompanying manual. The manual includes sheet music, explanations of musical terms, and guidance on effective practice routines, addressing prior findings that novices struggle with practice strategies (35,36). Participants practice for 30 minutes, five days per week (20 hours total), logging their sessions in diaries. Training progresses from single-hand exercises to bimanual coordination, incorporating scales, arpeggios, Hanon exercises, sight-reading familiar melodies, and metronome-based timing tasks linked to executive function. Videos feature music notation, a virtual keyboard, and overhead demonstrations, delivered weekly via WeTransfer to stagger content and monitor adherence.

PIANO-Cog was assessed in a feasibility RCT and the full-details of the protocol can be found in (34). Briefly, the primary aims of the study were to assess the acceptability of PIANO-Cog, as a remote, self-guided musical instruction programme for adults over age of 50, and the feasibility (recruitment, retention, and adherence) of conducting a fully-powered RCT to investigate the effects of PIANO-Cog on cognitive, motor and brain microstructural metrics in healthy ageing. The secondary aims were concerned with gaining effect size (ES) estimates of changes in cognition and brain microstructural measurements following training compared with a passive control. Cognitive assessment included EF tasks measuring performance in working memory capacity and updating, response suppression, and attention switching as well as verbal memory. Grey and white matter microstructure were assessed in regions of interest (ROIs) of cortical and subcortical networks involved in executive function, auditory, and motor processing that had previously been identified in the cross-sectional and longitudinal music studies discussed above (20–25,32,33) These were derived from the FreeSurfer v6 longitudinal pipeline (37) and included bilateral inferior frontal gyri (composed of pars triangularis and pars opercularis labels), precentral gyri, superior parietal lobules and transverse and superior temporal gyri as well as the basal ganglia, hippocampi and thalami.

White matter tracts of interest included the corpus callosum and corticospinal tract, which are important for bimanual coordination and motor control, as well as the arcuate fasciculus, which supports fronto–temporal interactions involved in language and auditory–motor integration. These tracts have previously been shown to differ between musicians and non-musicians, making them relevant targets for examining training-related microstructural change (27–29).

Brain microstructural measurements were derived from multi-shell high angular resolution diffusion imaging (HARDI) (38) data that were acquired with a maximum b-value of 6,000 s/mm² on a strong gradient (300mT/m) 3T Siemens Connectom scanner. These data were fit to the Neurite Orientation Density and Dispersion Imaging (NODDI) (39) and the Soma and Neurite Density Imaging (SANDI) (40) models to gain a number of complementary microstructural indices. NODDI and SANDI provide separate estimates of extracellular and intracellular diffusion signal fractions. Intracellular fractions are modelled with sticks (NODDI, SANDI) to capture neurites (axons and dendrites) and with spheres to capture soma shapes in grey matter (SANDI), yielding a number of complementary parameters to characterize grey and white matter microstructure. NODDI provides the intracellular signal fraction as an estimate of neurite density referred to as Neurite Density Index (NDI), the extracellular isotropic signal fraction (ISF) as an estimate of free water, and the orientation dispersion index (ODI) as an estimate of the neurite dispersion. SANDI yields parameters of apparent soma density (*fsoma*) and soma size (*Rsoma*) (intracellular fractions modelled by spheres), neurite density (*fneurite*) comparable to NDI from NODDI, and extracellular isotropic and anisotropic signal fraction (*fextra*). The composite hindered and restricted model of diffusion (CHARMED) (41) was applied to provide an additional estimate of axonal density with the restricted fraction (i.e. restricted fraction, FR) (modelled with cylinders). These microstructural measurements were employed to explore the biophysical properties of training-induced plasticity mechanisms in brain network regions known to be involved in music processing.

## 1 Method

Ethical approval was granted by Cardiff University School of Psychology Ethics Committee (EC.23.05.16.6801GRA). The trial was pre-registered with ISRCTN (https://doi.org/10.1186/ISRCTN11023869). The protocol for the PIANO-Cog trial has previously been published (34).

### 1.1 Participants

Forty-four participants (26 females) aged 51-80 years (*M =* 63.73*, SD =* 7.94) were recruited from Cardiff and the surrounding via poster advertisements, community groups for adults over 50, local active retirement Facebook groups, the Nextdoor app and the CUBRIC participant recruitment website. Participants were deemed eligible if they were >50 years old, fluent English speakers, had normal/corrected-to-normal vision and hearing, < 4 years of formal musical or dance training, and were not currently involved in any musical activities such as exercise-to-music classes, dance classes, choir singing, partaking in musicals or any musical instrument training, or any other cognitive training programmes. Participants were excluded if they had a neurological and/or psychiatric history which could affect learning such as dementia, stroke, traumatic brain injury or depression requiring hospitalization.

Participants with MRI contra-indications, such as pacemakers, stents or cochlear implants were not scanned but were still eligible for training and cognitive and motor testing.

### 1.2 Design

A two-arm, unblinded, randomised controlled feasibility trial was conducted. Testing sessions were held at baseline and at 8-week follow-up, which involved a 30-minute MRI scan and 2.5 hours of cognitive and motor assessment. Participants were pseudo-randomly allocated to either the piano or control group using R version 4.41, stratifying for sex (male/female) and age variables (50-64 years or 65+ years).

### 1.3 Procedure

Interested individuals who responded to study advertisements received a participant information sheet (PIS) via email. Participants were screened over an initial phone call for cognitive impairment using the Telephone Interview for Cognitive Status (TICS) (42), for <4 years previous music experience and for MR contraindications. Participants who were ineligible for scanning completed cognitive and motor tests only.

The study took place at the Cardiff University Brain research Imaging Centre (CUBRIC). At the first study visit, participants had the opportunity to read through the PIS again, and ask any questions before providing informed consent. Participants underwent a 30-minute scan on 3T Siemens Connectom system, and then 2.5 hours of behavioural testing. Breaks were offered as needed. Baseline assessments were conducted blind to group allocation as neither the participants nor the researcher administering baseline assessments were informed of group allocation until after the completion of all baseline assessments. Participants were then randomised to either a piano or control group. The passive control group were instructed to continue their usual activities during the 8-week period and to refrain from engaging in musical or cognitive training activities. Participants in the piano group received a keyboard and were instructed to practise for 30 minutes, five days per week for eight weeks, recording the duration and content of practice sessions in provided diaries. Researchers maintained weekly contact with participants in the piano group throughout the intervention period.

At follow-up, participants in the intervention group returned their keyboards and practice logs, and completed an evaluation survey of the training. Efforts were made to ensure that a blinded member of the research team conducted follow-up testing for both groups, but this was not always the case due to limited staffing. Following assessment, blinded research assistants recorded their estimate of each participant’s group allocation, their level of certainty (rated on a 10-point scale), and any reasons for possible unblinding.

#### 1.3.1 Intervention

Participants in the piano group received a training manual and weekly time-released tutorial videos and were asked to practice for 30 minutes per day, 5 days per week. Sixty-one key portable electric keyboards (MK-2000 by Gear4music) were provided for home use for the duration of the study. The training involved learning scales, arpeggios and songs and the videos were designed so that participants could follow along without any need for in-person instruction. Metronome-based rhythm exercises hypothesized to engage executive and sensorimotor networks were also included. Full details are provided elsewhere (34).

#### 1.3.2 Control group

The control group continued their usual activities with no intervention, and were asked not to take part in any music-based activities such as musical instrument classes, dance classes, choir singing or any cognitive training programmes. Control participants received the piano training manual and instruction videos on completion of follow-up visit as a thank-you for their participation and to encourage retention.

### 1.4 Outcome Measures

See (34) for full details on outcome measures.

#### 1.4.1 Screening measures

- **Telephone Interview for Cognitive Status (TICS)** (**42**): The TICS is a short, reliable measure of global cognition administered remotely, with scores ≤31/41 indicative of dementia.
- **Test of Premorbid Functioning (TOPF)** (**43**): The TOPF assesses verbal IQ by recording the number of correctly pronounced words from a list of 70 irregularly spelled words.
- **Patient Health Questionnaire-8 (PHQ-8)** (**44**): The PHQ-8 screens for current depression, with scores >10 indicating possible major depression; the self-harm item was omitted to minimize participant distress.

#### 1.4.2 Primary outcome measures

Our primary goal was to assess feasibility in terms of recruitment, retention, adherence rates and intervention acceptability. We used a traffic light system to interpret the results: green indicated that all feasibility rates were ≥70% and the trial was considered successful; amber indicated that no rates were <40% but at least one <70%, suggesting that the project required review and design modifications; and red indicated that at least one rate was <40%, suggesting that a future RCT would not be feasible.

- Recruitment is measured by the percentage of participants who were deemed eligible and who consented to taking part out of those who received the PIS. Reasons for ineligibility and declining to take part are reported in section 2.2.1.
- Retention is measured by the percentage of participants who completed both baseline and follow-up testing sessions. Reasons for withdrawal are reported in section 2.2.2.
- Adherence to the training was measured as frequency and duration of practice recorded in each participant’s practice diary, and by the percentage of training videos downloaded by each participant to track the feasibility of the technology used.
- Acceptability is measured using a self-report questionnaire, consisting of 27 Likert-scale items assessing quality, difficulty level and content of the training, with four questions providing opportunity for qualitative feedback.

#### 1.4.3 Secondary outcome measures

Secondary outcome measures were chosen to assess fluid abilities, executive functions, and verbal memory, which are most affected in ageing. Assessments targeted the three core EF: inhibitory control, attention switching, and working memory/updating, alongside processing speed and verbal memory. Near-transfer effects to music listening abilities were measured with the micro-PROMS (45). Brief descriptions are provided below and full details of all outcome measures are provided elsewhere (34).

- **Digit-Symbol Test (WAIS-III)** (**46**): Measures processing speed by recording the number of correct symbol-number pairings completed within 90 seconds.
- **Go/No-Go Test:** Assesses response inhibition by measuring accuracy and reaction time when participants respond to one stimulus but suppress responses to another.
- **Digit Span (Forward and Backward)** (**46**): Measures working memory by having participants recall sequences of numbers forwards or backwards.
- **N-Back:** Assesses updating of working memory through identification of stimuli repeated N steps earlier, including dual-task blocks for letters and spatial locations.
- **Trail-Making Test (TMT)** (**47**): Measures visual attention (Part A) and attention-switching (Part B) by recording completion times joining numbers or alternating numbers and letters.
- **D-KEFS Verbal Fluency** (**48**): Assesses word generation through letter fluency, category fluency, and category switching tasks, each with a 60-second limit.
- **California Verbal Learning Test II (CVLT-II)** (**49**): Measures verbal learning and memory across five trials with immediate and delayed recall, including cued and free recall formats.
- **Profile of Music Perception Skills (micro-PROMS)** (**45**): Objectively measures perceptual musical skills via short sequences testing pitch, rhythm, tempo, melody, tuning, and timbre (45).

### 1.5 MRI acquisition

MRI data were acquired on a 3T Siemens Connectom scanner (Siemens Healthcare, Erlangen, Germany) with strong magnetic gradients (300mT/m) at Cardiff University Brain Research Imaging Centre (CUBRIC).

T1-weighted (T1w) images were obtained using a magnetization-prepared 180° radio-frequency pulse with rapid gradient-echo (MPRAGE) sequence, with the following parameters: repetition time (TR) = 2,300 ms, echo time (TE) = 2 ms, inversion time (TI) = 857 ms, flip angle = 9°, field of view (FOV) = 256 × 256 × 192 mm, matrix size = 256 × 256 × 192, isotropic resolution = 1 × 1 × 1 mm³, phase-encoding direction = anterior-to-posterior (AP), and in-plane acceleration (GRAPPA) factor = 2, resulting in an acquisition time of 6 minutes.

The Connectom’s strong magnetic gradients allow for stronger diffusion weighting per unit time, which shortens the minimum echo time and thus improves signal to noise ratio, which is especially important at higher b-values, and increases sensitivity to small water displacement (50,51). Multi-shell High Angular Resolution Diffusion Imaging (ms-HARDI) (38) data were acquired using a single-shot spin-echo echo-planar imaging sequence with TR = 3,000 ms, TE = 59 ms, FOV = 220 × 200 mm, matrix size = 110 × 110 × 66, isotropic resolution = 2 mm³, gradient pulse duration (δ) = 7 ms, and gradient separation (Δ) = 24 ms, in the AP phase-encoding direction with GRAPPA factor = 2. Diffusion-weighted images were collected at b-values of 200 s/mm² (20 directions), 500 s/mm² (20 directions), 1,200 s/mm² (30 directions), 2,400 s/mm² (61 directions), 4,000 s/mm² (61 directions), and 6,000 s/mm² (61 directions). Fifteen non-diffusion-weighted (b = 0 s/mm²) images were also acquired: two at the beginning and 11 interspersed at the 33rd volume and every 20th volume thereafter in the AP direction, plus two images in the posterior-to-anterior (PA) direction. The total HARDI acquisition time was 18 minutes.

### 1.6 Image Processing

Data were hosted on XNAT (xnat.org), and were organised into standard Brain Imaging Data Structure (BIDS; https://bids.neuroimaging.io/index.html) format before processing.

#### 1.6.1 T1-weighted image preprocessing

The longitudinal pipeline from FreeSurfer (52) was utilised to segment cortical and subcortical ROIs according to the Desikan-Killiany atlas (53). This pipeline increases sensitivity to subtle longitudinal changes while reducing intra-subject variability in segmentation and surface reconstruction by creating an unbiased within-subject template from baseline and follow-up sessions for each participant, which improves consistency of volume estimates across sessions (54).

This pipeline includes removal of non-brain tissue, affine registration to Talairach space, intensity normalisation to correct for B1 bias field inhomogeneities, automated tissue classification, tessellation of the grey–white matter boundary, topology correction, and surface deformation guided by intensity gradients (37,55).

The volumetric segmentation files (.mgz) were converted to Nifti format using FreeSurfer, and then binary ROI masks were created from T1-weighted longitudinal run images using FSL (56).

#### 1.6.2 Diffusion-weighted Image Processing

An in-house modular pipeline, previously applied in the WAND study (57), was used to preprocess the multi-shell diffusion-weighted MRI data. The steps involved corrections for: thermal noise, signal drift, susceptibility-induced distortions, motion and eddy current-induced distortions, gradient non-linearity, and Gibbs ringing artefacts.

Brain extraction was performed by masking the first non-diffusion weighted image from each phase-encoding direction (A>P and P>A) to exclude all non-brain data using the FMRIB Software Library (FSL version 6.0.3 (56) brain extraction tool (58). A Marchenko-Pastur principal component analysis (MP-PCA)-based approach (59–61) in MRtrix3 (62) was used for diffusion noise level estimation and denoising. Then, within-image intensity drift was corrected by fitting the diffusion-weighted MRI data to temporally interspersed b0 (non-diffusion-weighted) images using in-house code in MATLAB R2017b (MathWorks Inc. Natick, Massachusetts, USA). Outlier detection was performed using Slicewise OutLIer Detection (SOLID; (63) with lower and upper thresholds of 3.5 and 10 respectively, using a modified Z-score and a variance-based intensity metric. FSL’s topup (64,65) was used to estimate the susceptibility-induced off-resonance field from the non-diffusion encoded (b0) data collected in opposing (A>P and P>A) phase-encoding directions and corrected, along with eddy current-induced distortions and subject movements using FSL’s eddy tool (66).

In-house code was used to correct for gradient non-linearity distortions in MATLAB R2017b (MathWorks Inc. Natick, Massachusetts, USA). Finally, MRtrix3 (62) was used to correct for Gibbs ringing using the method of local subvoxel-shifts proposed by (67) After preprocessing, MRtrix3 (62) was used to calculate the fibre orientation distribution function using multi-shell multi-tissue constrained spherical deconvolution (68).

#### 1.6.3 Diffusion Tensor Imaging (DTI)

DTI metrics were estimated using a nonlinear least-squares tensor fitting procedure implemented with an in-house MATLAB pipeline using b-values <2500 s/mm². Voxel-wise maps of fractional anisotropy (FA) and mean diffusivity (MD) were derived. FA reflects directional coherence of diffusion, while MD reflects overall diffusivity.

#### 1.6.4 Application of microstructure models (NODDI, SANDI and CHARMED)

NODDI (39) was applied to b-values 0, 500, 1,200, 2,400 s/mm², using the Microstructure Diffusion Toolbox (https://github.com/robbert-harms/MDT) to extract estimates of intracellular signal fraction (ICSF), orientation dispersion index (ODI) and extracellular signal fraction (ECSF).

All b-values (200 s/mm², 500 s/mm², 1,200 s/mm², 2,400 s/mm², 4,000 s/mm² and 6,000 s/mm²) were used for SANDI and CHARMED fitting. For SANDI-fitting, noise maps were generated for each subject’s diffusion-weighted images using a MP-PCA-based method (59–61) in MRtrix3 (62). These noise maps were then used to fit the SANDI model to the pre-processed multi-shell diffusion data using the SANDI Toolbox (https://github.com/palombom/SANDI-Matlab-Toolbox-Latest-Release) using default settings. The toolbox produced intra-soma signal fraction (*fsoma*), intra-neurite (*fneurite*) and extracellular water (*fextra*) signal fractions, apparent soma size (*Rsoma*), and extracellular diffusivity (*De*).

CHARMED (41) analysis was performed using in-house code implemented in MATLAB R2017b. A standard CHARMED model was first fitted, incorporating information derived from multi-shell, multi-tissue constrained spherical deconvolution fibre orientation distributions to determine the number of compartments per voxel and to constrain their orientations. The parameters estimated during this initial fitting stage were subsequently used to compute restricted fraction maps, which quantify the total amount of restricted diffusion within each voxel.

#### 1.6.5 Co-registration, Partial Volume Correction and Extraction of Microstructural Metrics from Grey Matter ROIs

Registration matrices were created by registering FA maps to the longitudinal T1-weighted structural images using Advanced Normalization Tools (ANTS) (69). Registration was performed in three dimensions using a multi-stage linear framework consisting of rigid and affine transformations. Mutual information was used as the similarity metric at each stage, with a multi-resolution optimization scheme (four resolution levels, decreasing smoothing and shrink factors). Histogram matching and intensity winsorization (0.5–99.5%) were applied to improve robustness. The resulting matrices were inversed (T1-FA) and used to transform ROIs to microstructural diffusion maps from NODDI and SANDI, as well as FR and MD.

Subcortical ROIs (basal ganglia structures, hippocampi and thalami) were eroded by one voxel to minimise partial volume effects due to proximity with CSF-filled ventricles. For cortical ROIs, partial volume correction was applied by acquiring CSF maps of the T1 longitudinal runs using FreeSurfer (37), extracting the mean CSF values from each ROI and including those values as covariates for each ROI.

Means, standard deviations and 25^th^, 50^th^ and 75^th^ percentile values of each microstructural index from DTI (FA, MD), the restricted fraction index (FR) from CHARMED (41), the neurite density index (NDI), extracellular isotropic signal fraction (ISF) and orientation dispersion index (ODI) from NODDI (39) and the intra-soma signal fraction, soma size, intra-neurite signal fraction, extracellular signal fraction and extracellular diffusivity from SANDI (40) were extracted for each ROI using *fslmaths* from FSL (56).

#### 1.6.6 Tractometry

White matter TOIs – the corpus callosum, bilateral arcuate fasciculus, corticospinal tract and fornix - were reconstructed using TractSeg v2.9 (70), Python v 3.8.20. All TOIs were visually inspected using Fibernavigator (https://scilus.github.io/fibernavigator/). Sparse tracts and tracts that failed to reconstruct were recorded and are reported in subsection 2.3.3.

Tractometry was used to obtain 100 along-tract FA statistics for the corpus callosum, bilateral arcuate fasciculus and corticospinal tract. TractSeg applies machine learning to create tract masks as well as tract orientation maps and start/end region masks for each tract which allow for the generation of accurate bundle-specific tractograms. TractSeg is fully automated approach to white matter tract segmentation which segments white matter tracts in fields of fiber orientation distribution (FOD) peaks. FOD peaks indicate the main directions of fibers in each voxel. All streamlines leaving the tract mask and not ending in the start/end masks are discarded, resulting in one tractogram for each tract. TractSeg code was accessed on the openly available (https://github.com/MIC-DKFZ/TractSeg) with pretrained weights. The super resolution setting was used which outputs the image at 1.25m.

Tractometry was not performed for the fornix because tractseg did not reliably reconstruct this tract due to its small size. Median, and interquartile ranges (IQRs) for NODDI, SANDI and FR metrics were extracted from the fornix and other tracts by registering the maps to the binary bundle segmentation masks generated by TractSeg.

#### 1.6.7 Statistical Analysis

Statistical analyses were performed in R version 4.5.2 in R-studio hosted on Posit Cloud (https://posit.cloud/). As the primary goal of this study was to evaluate feasibility, no formal power analysis was conducted in line Consolidated Standards of Reporting Trials (CONSORT) statement extended for pilot and feasibility studies (71).

##### 1.6.7.1 Feasibility Outcomes

Feasibility was assessed using predefined recruitment, retention, and adherence criteria. Recruitment was the proportion of eligible participants who consented, retention was the proportion completing post-intervention assessment, and adherence was the proportion of prescribed training completed. Outcomes were classified using a traffic-light system: green (≥70% for all metrics) indicated success, amber (≥40% but <70% for at least one) indicated design review needed, and red (<40% for any metric) indicated the intervention or future RCT was not feasible.

##### 1.6.7.2 Baseline Group Comparisons

Kruskal-Wallis tests were performed to test for baseline group differences in age, TICS, TOPF and micro-PROMS scores. Normality was assessed using histograms and Shapiro-Wilk tests with alpha level of .05.

##### 1.6.7.3 Behavioural Outcomes

Changes in cognitive and motor performance were calculated as post-intervention minus pre-intervention scores, such that positive values reflected improvement over time. For tasks with multiple conditions (e.g., verbal fluency, N-back), change scores were computed separately for each condition. Group differences in change scores were summarised descriptively and visualised using forest plots of standardised mean differences (Hedge’s g), line plots of raw pre- and post-intervention scores, and violin plots of change distributions. Given the exploratory nature of these analyses, emphasis was placed on effect sizes and patterns of change rather than statistical significance.

##### 1.6.7.4 Microstructural outcomes

Microstructural diffusion metrics were extracted on a voxel-wise basis and summarised within regions of interest. Change in microstructural measures was calculated as post-intervention minus pre-intervention values. Group differences in microstructural change were explored descriptively using standardised mean differences and visual inspection of spatial maps and summary plots. These analyses were considered hypothesis-generating and interpreted cautiously.

## 2 Results

### 2.1 Sample Information

Demographic variables for all participants who attended both baseline and 8-week follow-up sessions are presented in Table 1, along with the subset of participants who were eligible for MRI scanning and included in the microstructure analysis. Of the 44 participants who attended the baseline session, 18 did not undergo MRI scanning for the following reasons: 8 participants with a shoulder/hip width exceeding 54 cm, precluding scanner entry (bore diameter: 56cm), 3 participants with medical stents, 3 who were claustrophobic, 2 prone to migraines; 1 with a magnetic prosthetic ear; and 1 who had recently undergone a dental procedure. Of the 26 participants scanned at baseline, 18 were also scanned at follow-up.

**Table 1:** Group demographic characteristics for participants that attended baseline and follow-up sessions.

| Group | Age (years) | Sex (M/F) | Education (years) | TICS | TOPF | Employment<br>(FT/PT/Retired) | microPROMS |
| --- | --- | --- | --- | --- | --- | --- | --- |
| All participants who attended baseline and follow-up testing sessions |  |  |  |  |  |  |  |
| piano | 63.5 (9.4)<br>[52–79] | 6/9 | 16.1 (3.8)<br>[8–22] | 30.8 (2.8)<br>[24–34] | 63.9 (3.7)<br>[56–69] | 4/3/8 | 10.0 (2.7)<br>[5–16] |
| control | 65.7 (5.0)<br>[57–74] | 5/13 | 16.3 (3.8)<br>[9–25] | 29.8 (2.8)<br>[25–36] | 58.2 (6.0)<br>[48–66] | 2/4/12 | 8.9 (1.5)<br>[6–12] |
| Participants who attended baseline and follow-up testing sessions who underwent MRI scanning |  |  |  |  |  |  |  |
| piano | 61.0 (9.6)<br>[52–79] | 2/4 | 16.0 (5.1)<br>[8–22] | 31.5 (2.3)<br>[29–34] | 65.0 (2.2)<br>[62–68] | 1/3/2 | 8.3 (2.0)<br>[5–10] |
| control | 65.9 (5.7)<br>[57–74] | 1/8 | 17.6 (3.8)<br>[13–25] | 28.9 (2.0)<br>[26–31] | 57.4 (4.7)<br>[53–66] | 1/3/5 | 8.7 (1.4)<br>[6–10] |

Continuous variables are mean (SD) [Min–Max]. Categorical variables are counts (see labels). TICS, Telephone Interview for Cognitive Status, TOPF, Test of Premorbid Functioning, microPROMS, Profile of Music Perception Skills

Kruskal-Wallis tests revealed no statistically significant differences between piano and control groups at baseline in age, global cognitive ability (TICS), baseline music perception abilities (micro-PROMS) or any cognitive measures (see Supplementary material). However, there was a statistically significant difference in TOPF scores at baseline (*W* = 79.00, *p =* .001) with the piano group having higher baseline scores (median = 65) than the control group (median = 57.50).

### 2.2 Primary Outcomes: Feasibility

#### 2.2.1 Recruitment

Of the 113 individuals who responded to our study advertisements and received a participant information sheet, 48 (42.5%) were eligible to participate, 16 (14.2%) were ineligible and 49 did not respond after receiving a PIS and were therefore not screened.

Out of the 48 participants who were eligible, 44 consented to participate (91.7%), indicating a green traffic light rating *(Figure 1)*. Reasons for declining to participate were: travel distance, family bereavement, lack of availability for testing session and loss of contact.

**Figure 1:**
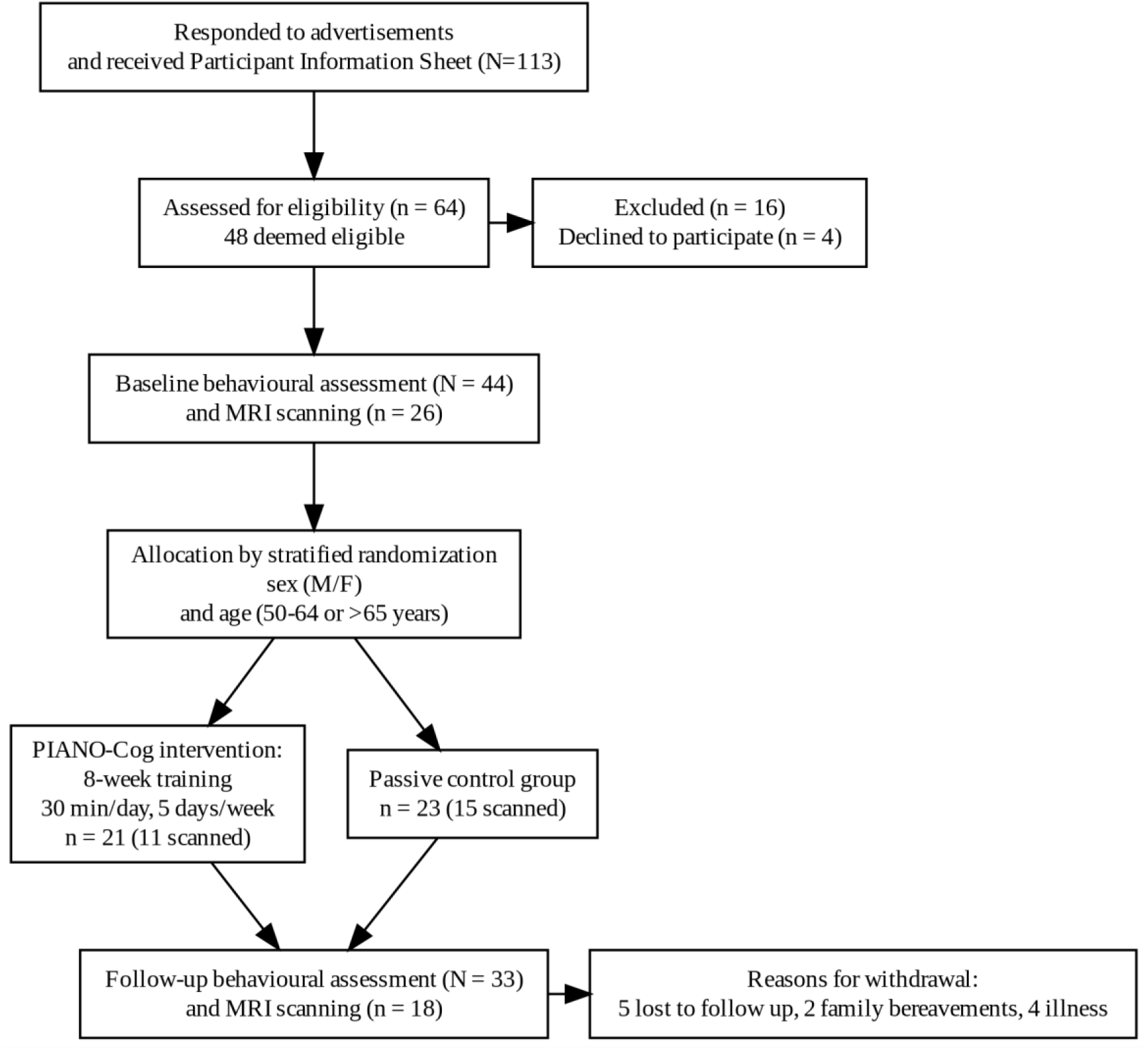
CONSORT flow diagram of feasibility RCT.

#### 2.2.2 Retention

33 out of 44 participants attended follow-up. 6 dropped out of the piano group (2 due to injury/illness, 1 family bereavement and 3 lost to follow-up) and 5 dropped out of the control group (1 family bereavement, 2 illness, 2 lost to follow-up).

#### 2.2.3 Adherence

The intervention group were required to download and watch the new training video every week and to practise for 30 minutes, 5 days per week (2.5 hours per week, or 20 hours across 40 days total). Table 2 demonstrates that that frequency and duration measures were rated as green (>70%) across the participants assigned to piano training according to the predetermined traffic light system.

**Table 2:**
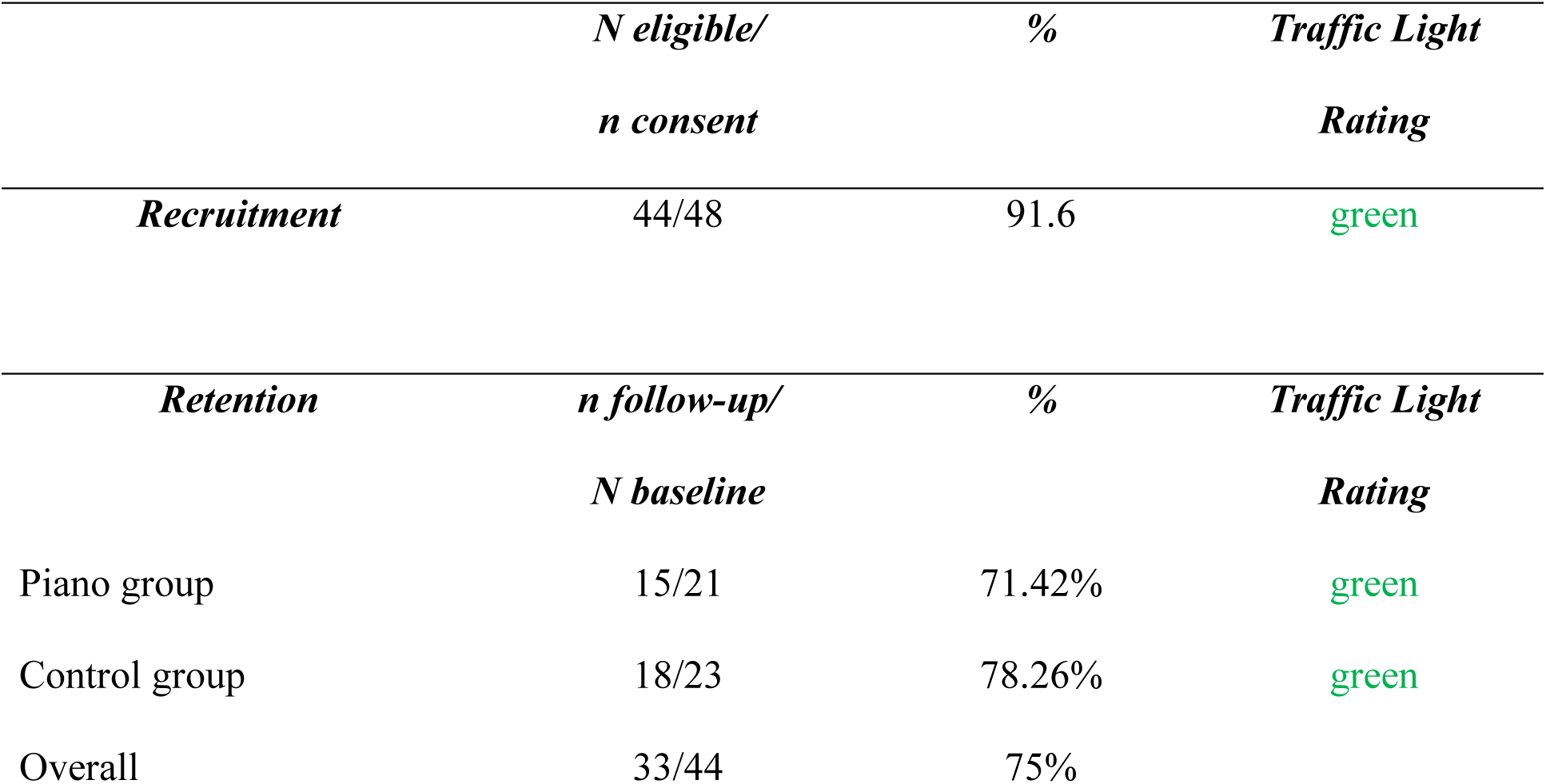
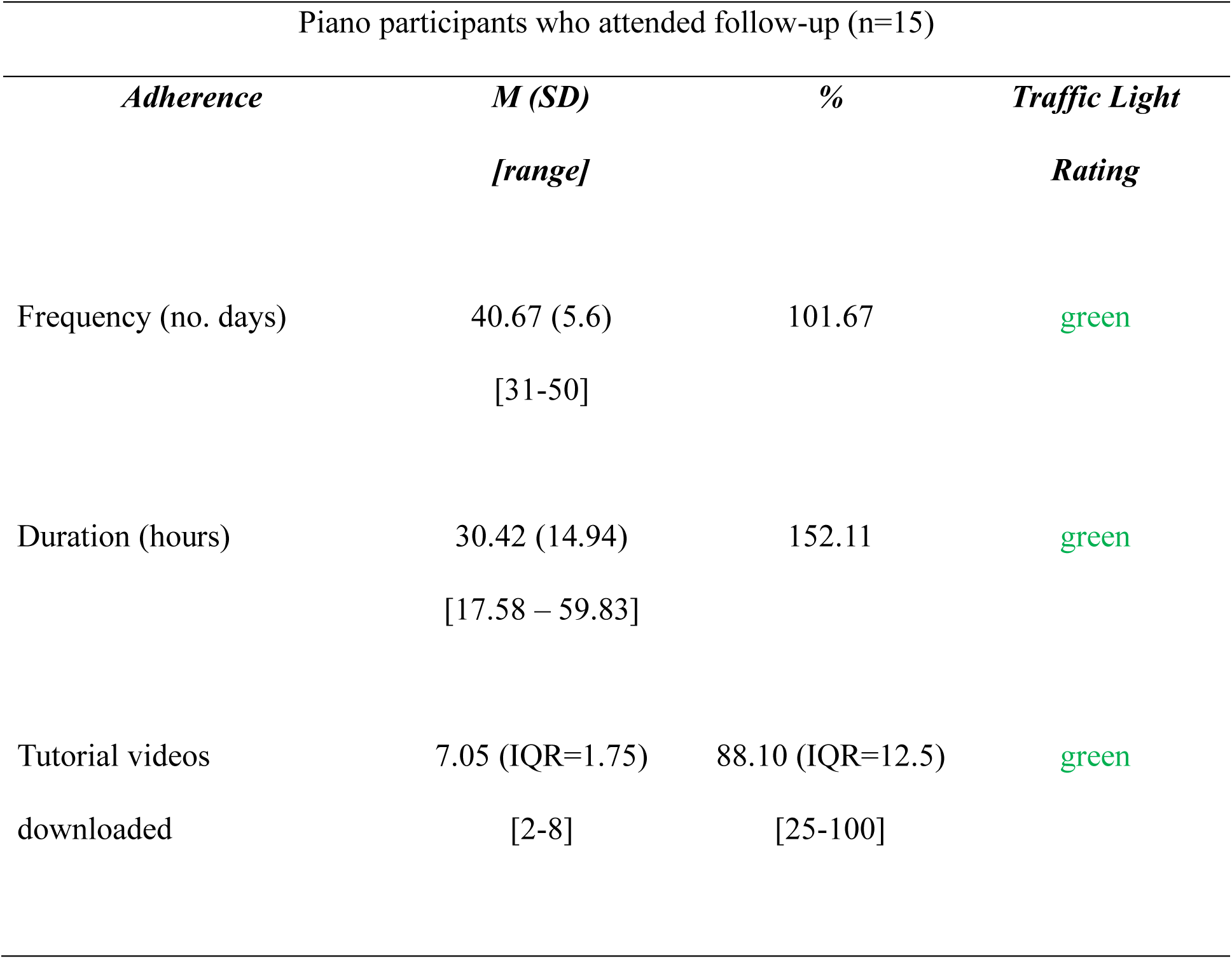
Descriptive statistics for feasibility outcome measures (recruitment, retention and adherence for the piano intervention.

2.2.4 Acceptability:

*Figure 2* presents percentage scores of Likert item responses from PIANO-Cog evaluation survey.

**Figure 2:**
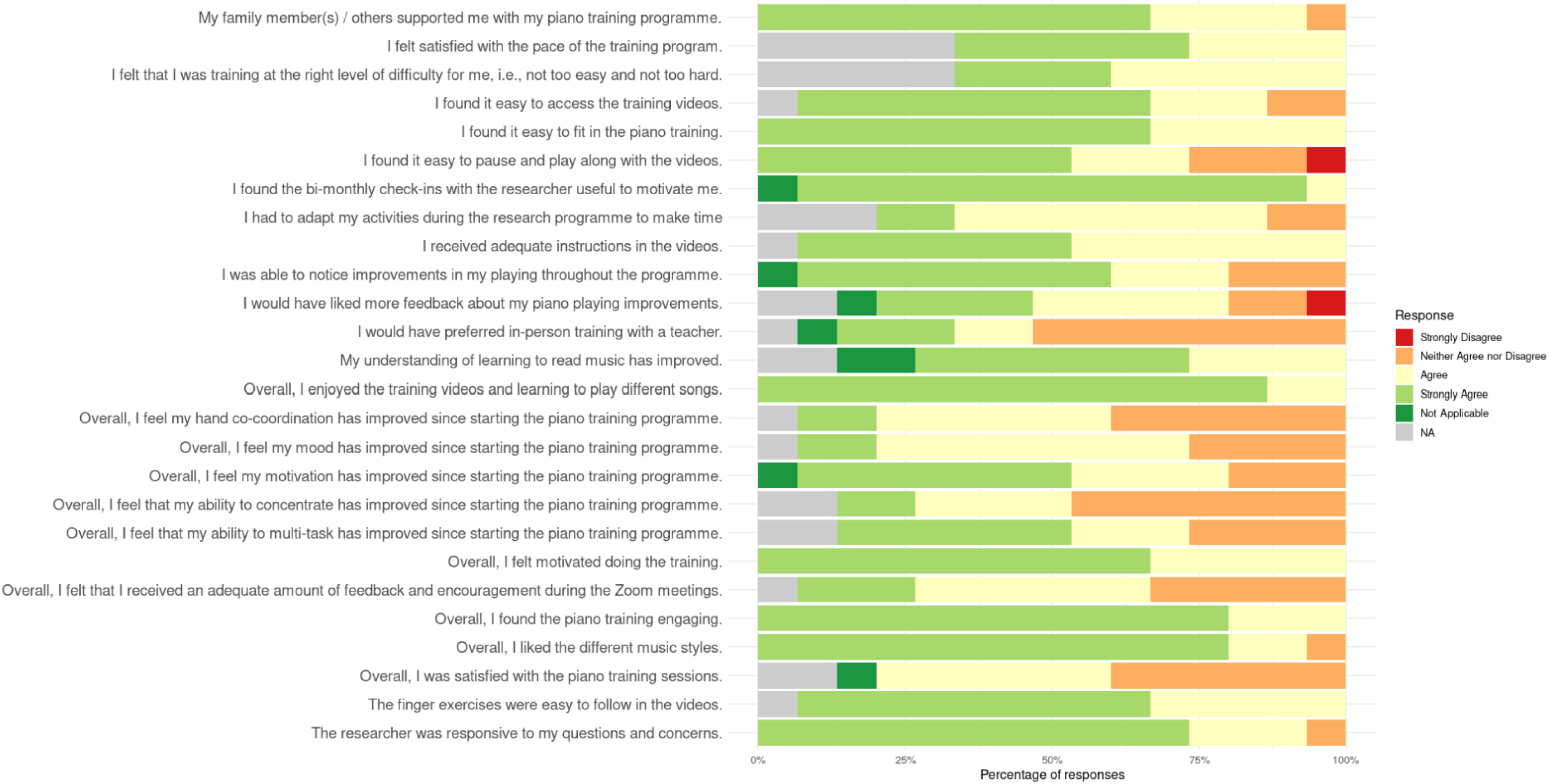
Responses on Likert items form Piano Evaluation Survey (n=15).

### 2.3 Secondary Outcomes

#### 2.3.1 Cognitive Outcome Measures

Pen and paper and oral tests are presented in *Figure 3*, and computerised tests, Go/No-Go and N-Back are described separately below. Letter fluency was the only test which showed a significant increase in Hedge’s G in the piano group compared to the control group, out of the verbal and paper-and-pen assessments.

**Figure 3:**
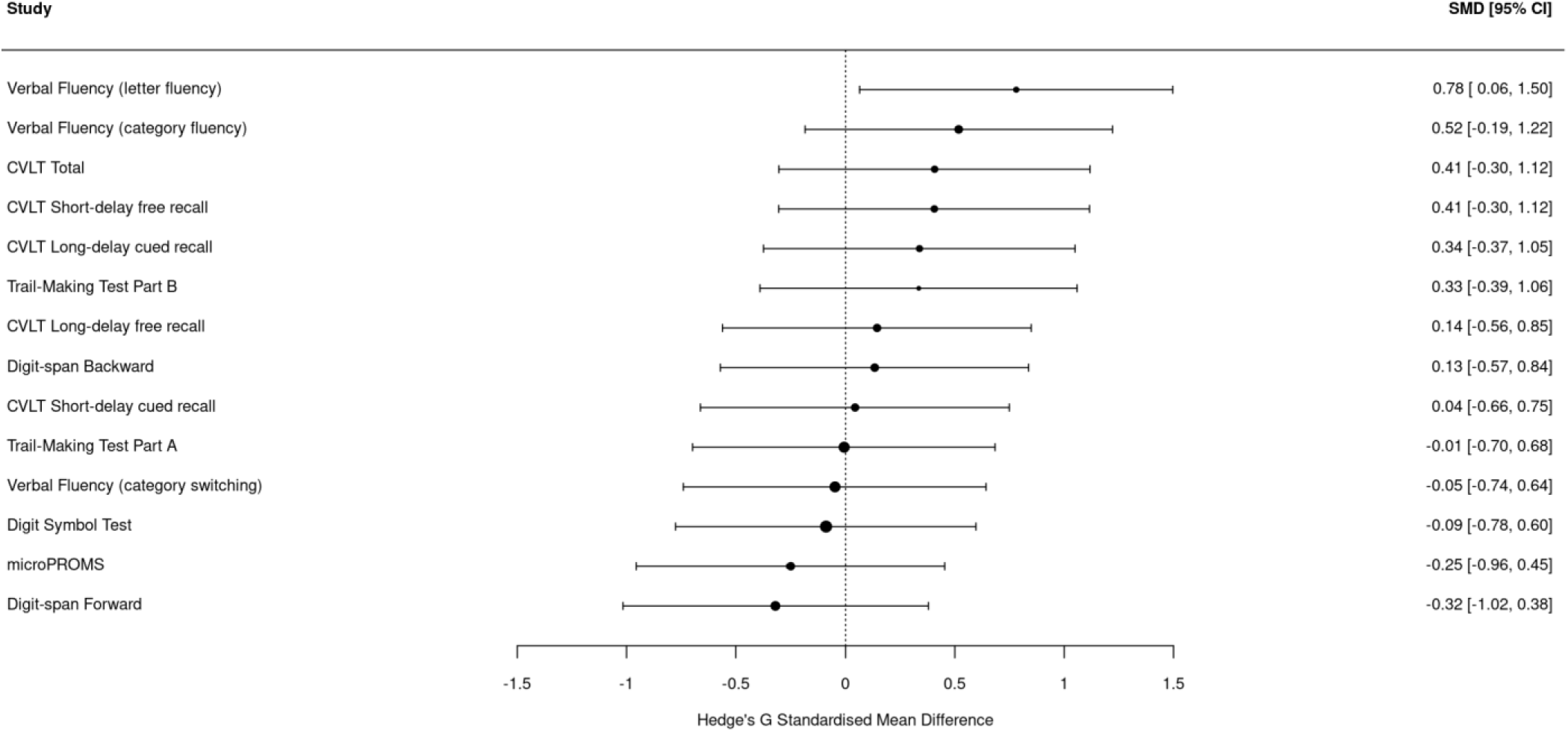
Forest plot presenting Hedge’s G Standarised Mean Difference in baseline-follow-up change scores for piano versus control group, with scores to the right and side suggesting greater improvement in piano group.

##### 2.3.1.1 Go/No-Go

Mean change from baseline to follow-up in the total number of hits, misses, correct passes and false alarms were compared between groups for the Go/No-Go analysis.

Three participants were removed from the analysis: the number of hits and misses for two participants (1 piano and 1 control) were identified as extreme outliers (a factor of 3 x IQR beyond the 25^th^ percentile); and 1 control participant was not included due to technical difficulties on the day of testing. This left 15 piano and 16 control participants included in the analyses. 17 trials were removed from the entire sample because they were <200ms, and therefore unreliable as true hits/false alarms.

A trend was identified where the piano group showed a greater increase in the number of hits and a decrease in the number of misses at follow-up compared to baseline, suggesting improved performance for the piano group *(Figure 4).* Line plots and violin plots are available in supplementary material. No change was seen in the number of false alarms or correct passes for either group. See descriptive statistics in Table 3.

**Figure 4:**
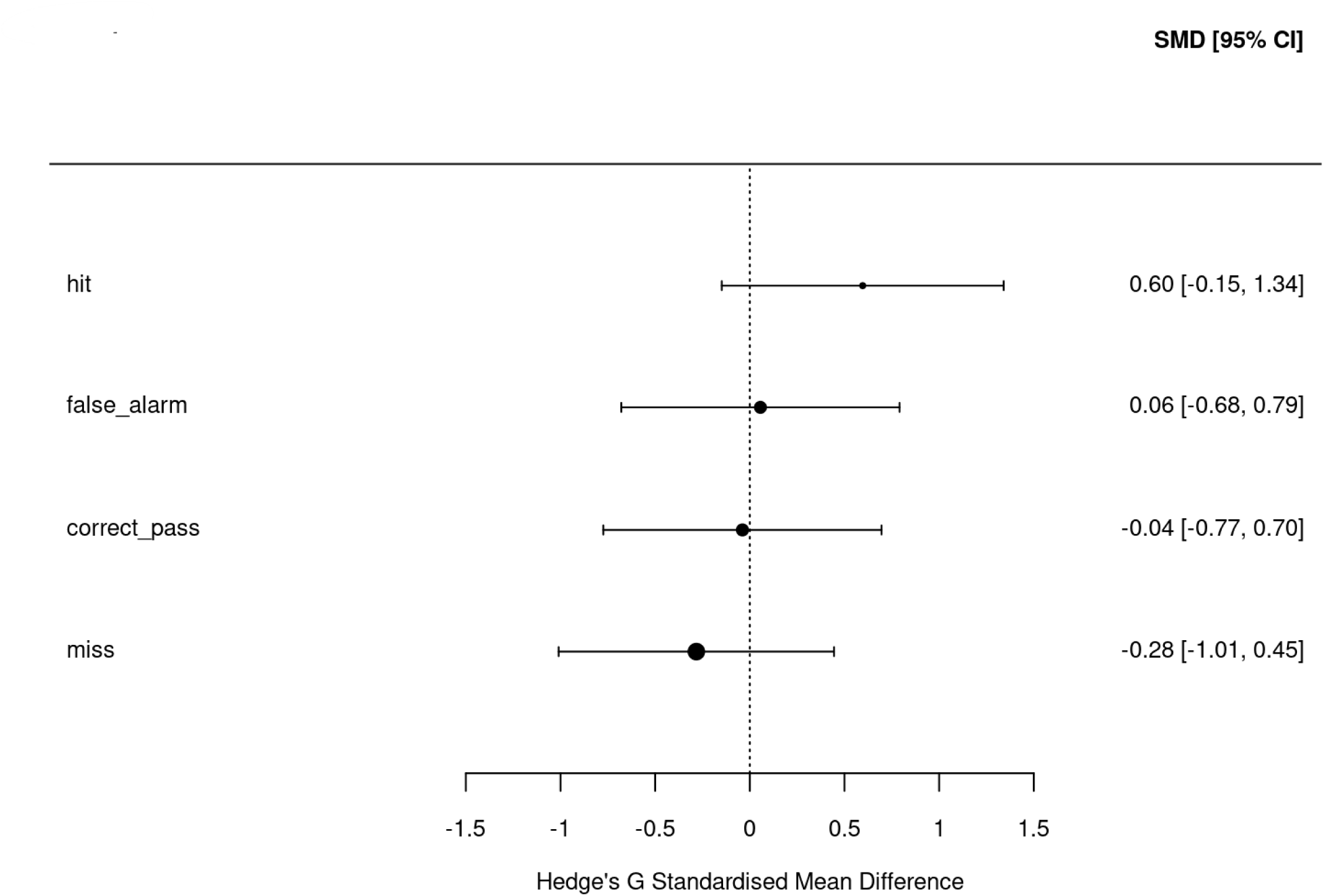
Forest plot of differences in change scores on Go/No-Go trials.

**Table 3:** Descriptive statistics for Go/No-Go trials for piano and control groups across sessions.

|  | *Piano* |  |  |  | *Control* |  |  |  |
| --- | --- | --- | --- | --- | --- | --- | --- | --- |
|  | *Mean (SD)* |  |  |  | *Mean (SD)* |  |  |  |
|  | *[range]* |  |  |  | *[range]* |  |  |  |
|  | *hits* | *misses* | *false alarms* | *correct passes* | *hits* | *misses* | *false alarms* | *correct passes* |
| *Baseline* | 254.80 (1.61) | 0.80 (1.08) | 6.80 (3.69) | 57.00 (3.70) | 255.19 (1.22) | 0.75 (1.13) | 9.50 (5.11) | 54.50 (5.11) |
|  | [252 - 256] | [0 – 3] | [2 - 17] | [47 – 62] | [253 - 256] | [0 - 3] | [2 – 20] | [44 - 62] |
| *Follow-up* | 255.33 (1.08) | 0.53 (0.63) | 5.27 (3.81) | 58.60 (3.87) | 255.00 (1.41) | 0.81 (1.38) | 7.75 (3.73) | 56.25 (3.73) |
|  | [254 – 256] | [0 – 2] | [2 – 13] | [51 – 62] | [251 - 256] | [0 - 5] | [2 - 18] | [46 - 62] |

##### 2.3.1.2 N-Back

Mean change in the number of correct responses on letter, square and dual-task trials for 1-and 2-back blocks was assessed and compared between groups. A trend was observed where piano participants showed a larger increase in performance on the dual 2-back condition and the 2-back letter condition, suggesting improvement in auditory working memory updating compared to the control group Figure 5. See Table 4 for descriptive statistics. Line plots and violin plots are presented in supplementary material

**Figure 5:**
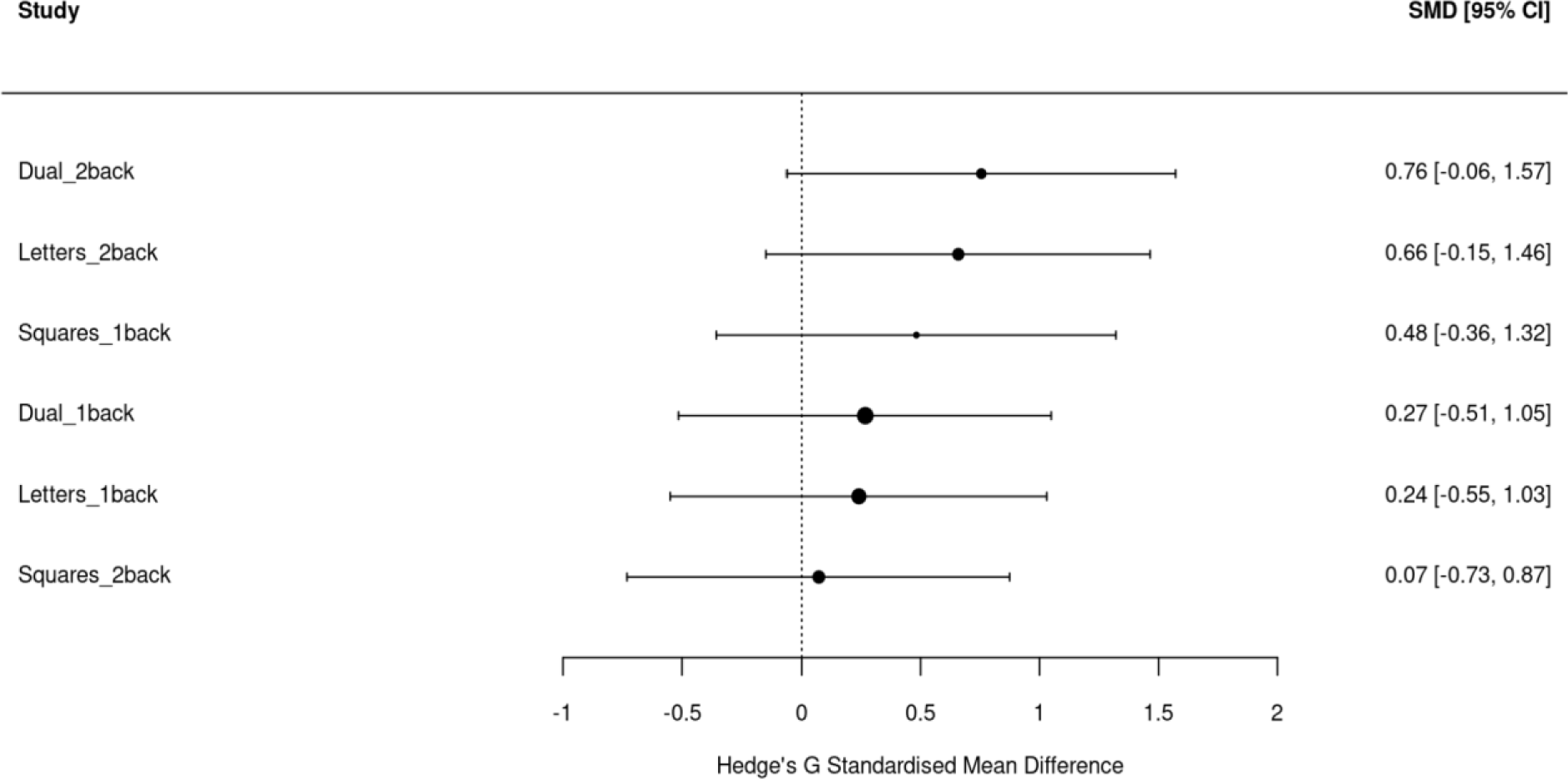
Forest plot of differences in change scores on N-Back trials.

**Table 4:** Descriptive statistics for N-back trials for piano and control groups across sessions.

|  |  | **Piano** |  |  | **Control** |  |  |
| --- | --- | --- | --- | --- | --- | --- | --- |
|  |  | *Mean (SD)* |  |  | *Mean (SD)* |  |  |
|  |  | *[range]* |  |  | *[range]* |  |  |
|  |  | *Letters* | *Squares* | *Dual-task* | *Letters* | *Squares* | *Dual-task* |
| *Baseline* | *1-back* | 19.60 (2.13) | 19.07 (2.55) | 18.53 (2.95) | 19.88 (1.36) | 18.94 (1.98) | 18.35 (2.47) |
|  |  | [14 - 21] | [13 – 21] | [12 – 21] | [16 – 21] | [15 – 21] | [13 – 21] |
|  | *2-back* | 17.93 (2.66) | 17.53 (1.73) | 14.60 (2.16) | 18.53 (2.53) | 17.18 (2.07) | 14.71 (2.49) |
|  |  | [14 - 22] | [15 – 20] | [11 – 19] | [13 – 22] | [14 – 21] | [10 – 20] |
| *Follow-up* | *1-back* | 19.64 (1.03) | 20.18 (0.99) | 18.91 (1.14) | 19.69 (1.85) | 19.00 (2.16) | 18.19 (3.17) |

|  | [17 - 21] | [19 - 21] | [17 - 21] | [15 - 21] | [15 - 21] | [11 - 21] |
| --- | --- | --- | --- | --- | --- | --- |
| *2-back* | 19.55 (1.37) | 17.36 (2.60) | 16.27 (2.41) | 18.50 (1.63) | 17.25 (1.95) | 14.75 (2.18) |
|  | [18 - 22] | [15 - 21] | [14 - 21] | [15 - 21] | [13 - 20] | [10 - 18] |

#### 2.3.2 Grey Matter ROIs

Cortical grey matter regions of interest included: (i) bilateral inferior frontal gyri (IFG), (ii) mid temporal gyri, (iii) transverstemporal gyri, (iv) banks of the superior temporal sulcus (STS), (v) superior parietal lobules, and (vi) primary motor cortices. Subcortical grey matter ROIs included (vii) bilateral structures of the basal ganglia (caudate nuclei, putamen and pallidum), (viii) hippocampi and (ix) thalami.

Forest plots of Hedges’s G standardized mean differences of group differences in change scores of microstructural measurements from baseline to follow-up (i.e. differences-in-differences) of all ROIs are shown in *Figure 6*. Exploratory microstructural analyses showed increases in *fsoma* and decreases in *fextra* in the left inferior frontal gyrus; increases in *fneurite* and NDI in the right inferior frontal gyrus; increased ODI and decreased FA in the left middle temporal gyrus; increased *Rsoma* and De in the right middle temporal gyrus; increased *Rsoma* and decreased *fneurite* in the right banks of the superior temporal sulcus; and increased ODI in the right precentral gyrus.

**Figure 6:**
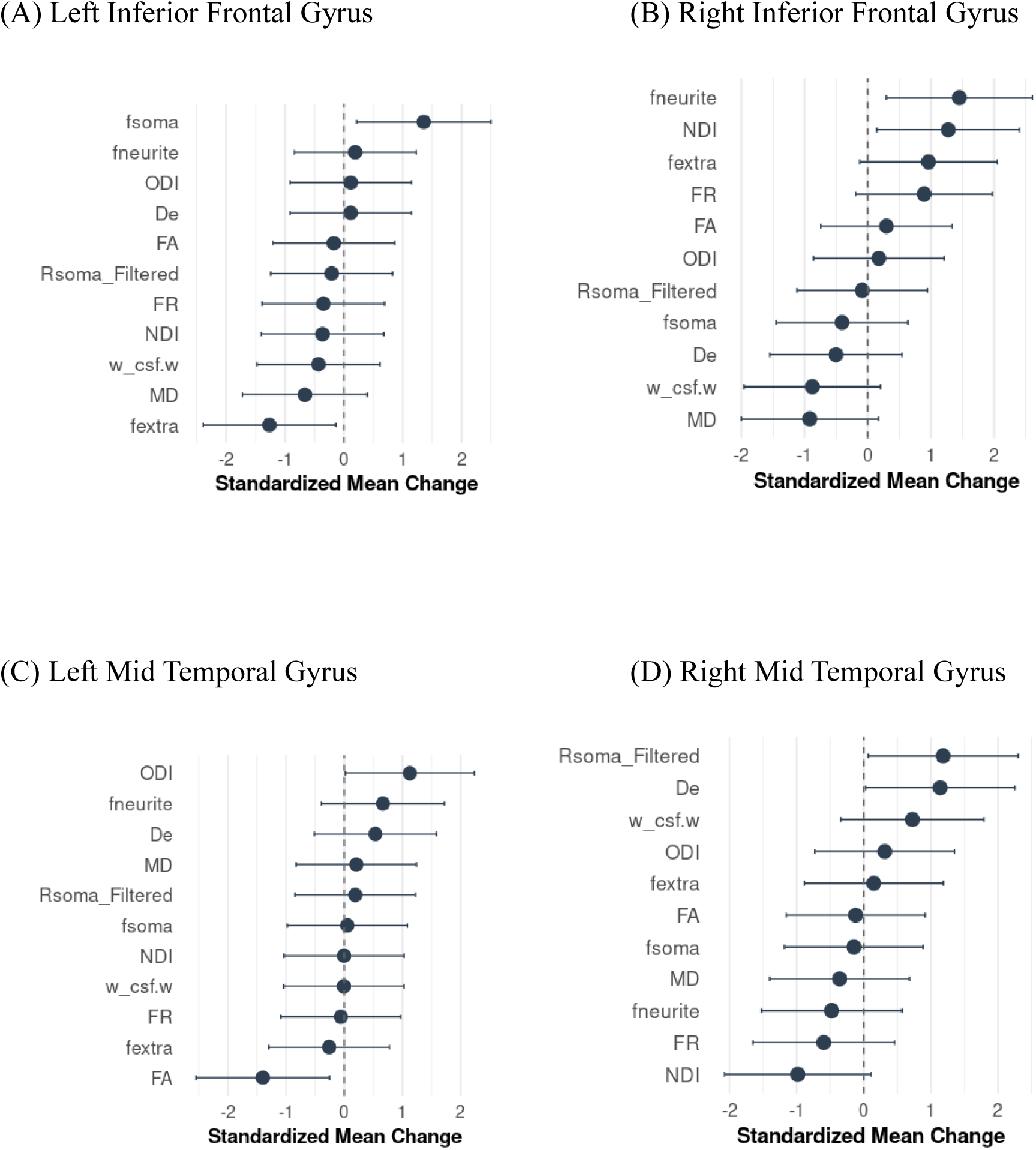
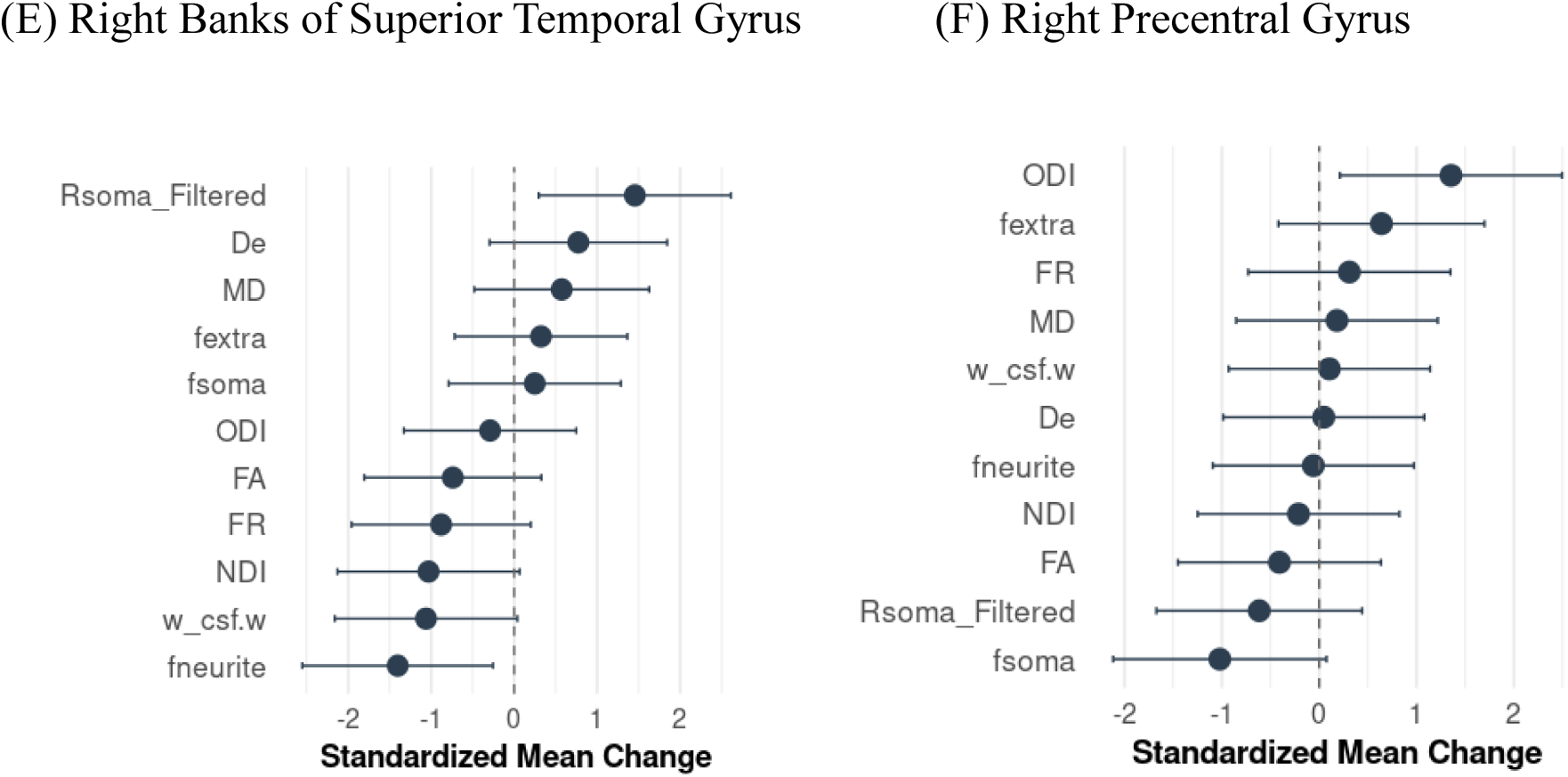
Forest plots presenting partial-volume corrected standardized mean group differences (Hedge’s G) in change scores for left (A) and right (B) inferior frontal gyri, left (C) and right (D) mid temporal gyri, (E) right banks of the superior temporal sulcus, (F) right precentral gyrus. Values to the right-hand side of the 0 line indicate a greater increase in the piano versus control group.

The following subsections detail specific microstructural measurement changes in these regions with line plots presenting raw data at baseline and follow-up, and violin plots showing group differences in value change (differences-in-differences). Means and SDs of baseline and follow-up are presented in Table 5

**Table 5:**
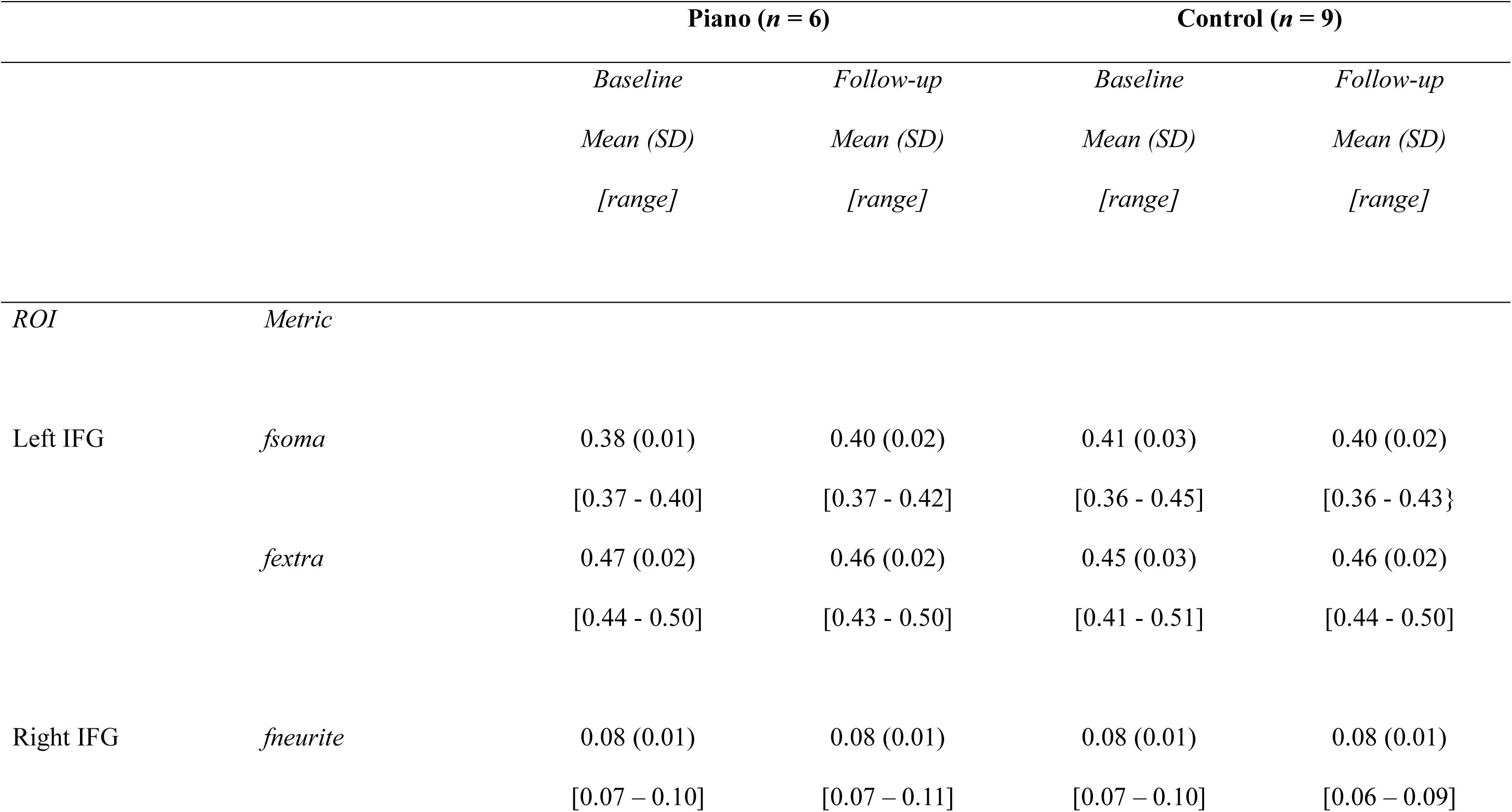
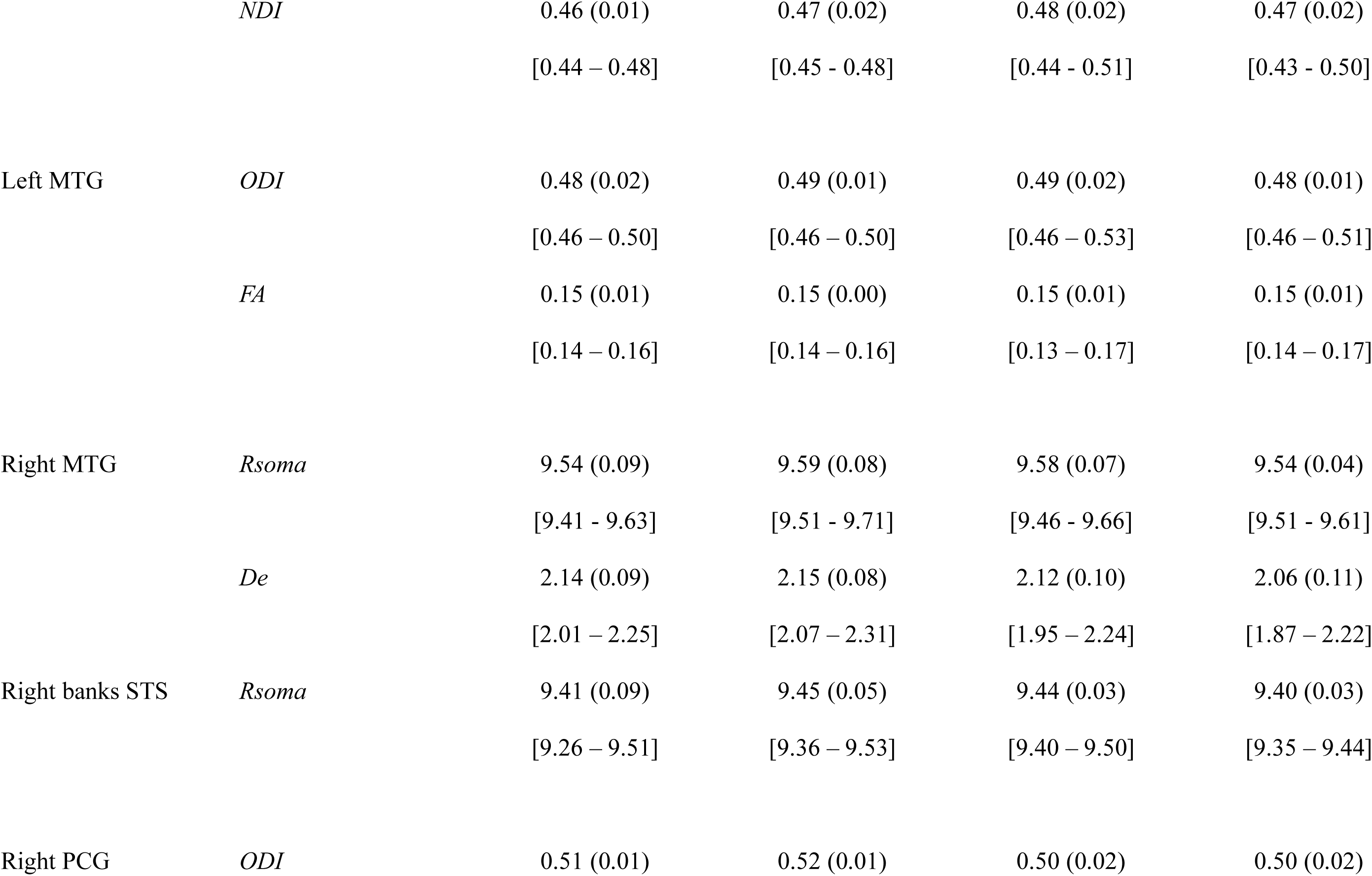
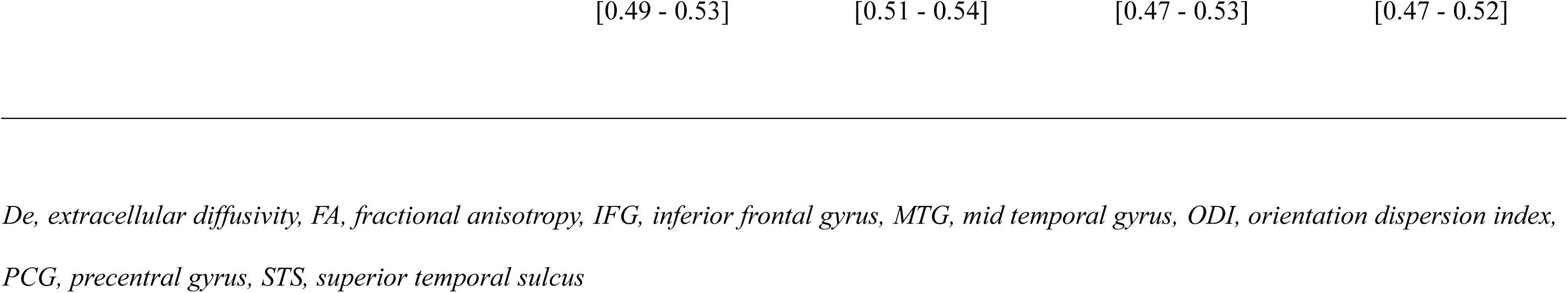
Descriptive statistics for microstructural metrics of grey matter ROIs for piano and control groups.

##### Inferior Frontal Gyri

An increase in *fsoma* and a decrease in *fextra* was observed for the piano group compared to the control group in the left IFG (Broca’s area), as demonstrated by the line plots and violin plots in Figure 7.

**Figure 7:**
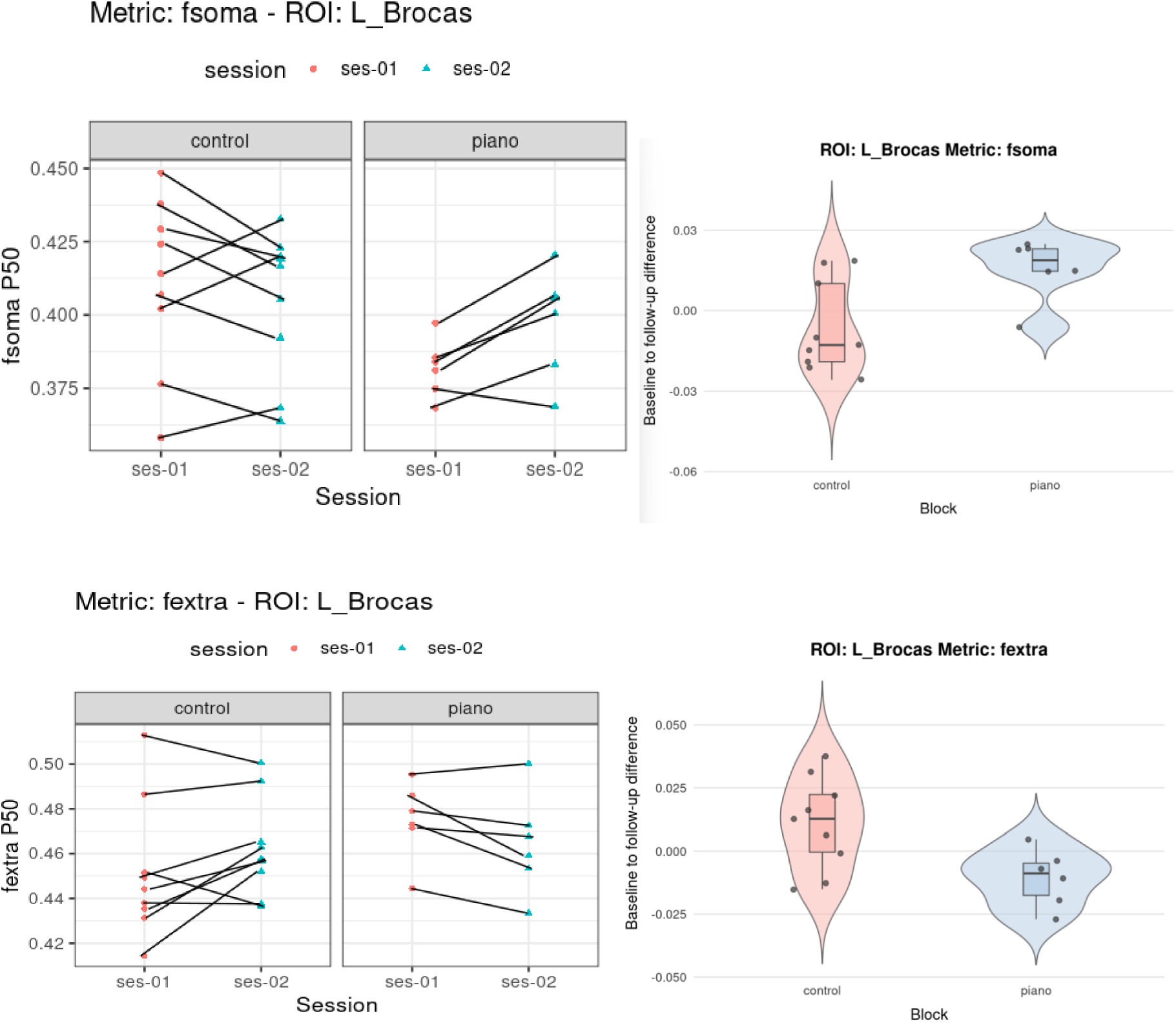
Line and violin plots demonstrating changes in fsoma and fextra in left inferior frontal gyrus in piano versus control group.

In contrast, the right IFG in piano participants displayed a slight increase or maintenance in *fneurite* from SANDI and NDI from NODDI measures of neurite density, whereas the control group tended to show a decrease over time (*Figure 8*).

**Figure 8:**
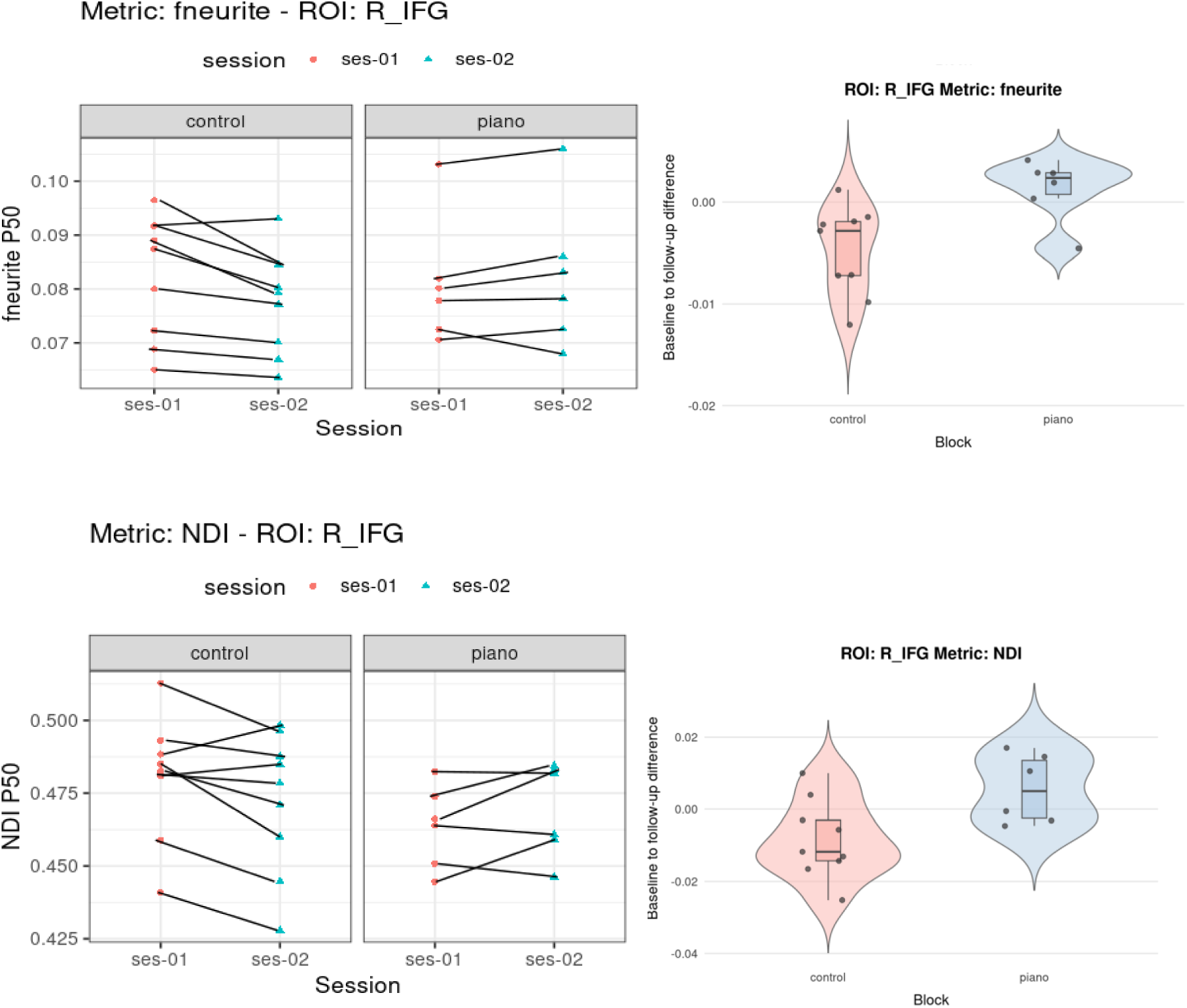
Line and violin plots demonstrating changes in NDI and fneurite in rightt inferior frontal gyrus in piano versus control group.

##### Mid Temporal Gyri

Neurite dispersion measured using the ODI appeared to remain stable for the piano group and showed a decrease in the control group in the left mid temporal gyrus accompanied by changes in FA *(Figure 9)*.

**Figure 9:**
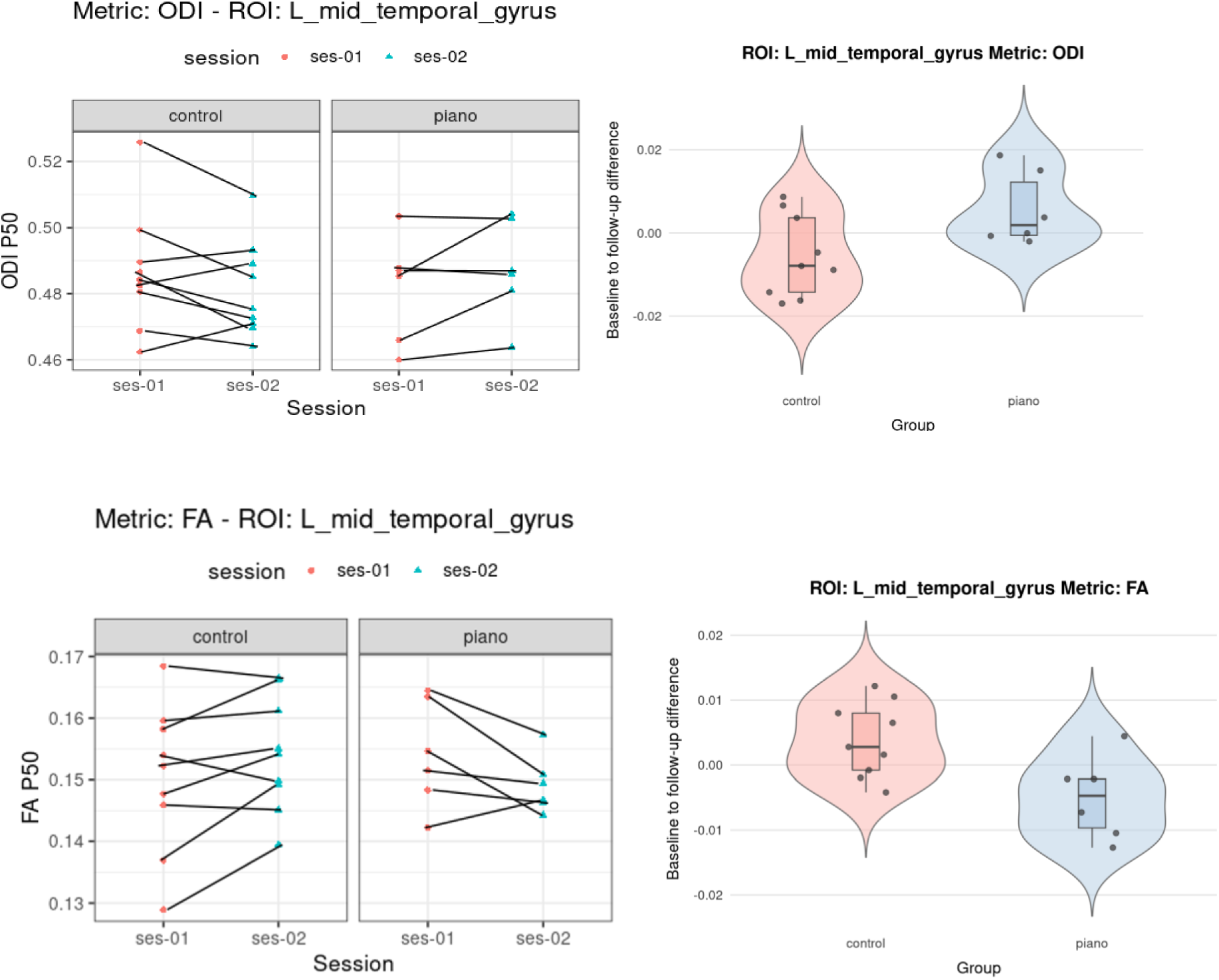
Line and violin plots demonstrating changes in ODI and FA in left mid temporal gyrus in piano versus control group.

Conversely, right mid temporal gyrus showed an increase in soma size for the piano group whereas it showed a decline for several control participants, accompanied by a decrease in extracellular diffusivity in the control group *(Figure 10)*.

**Figure 10:**
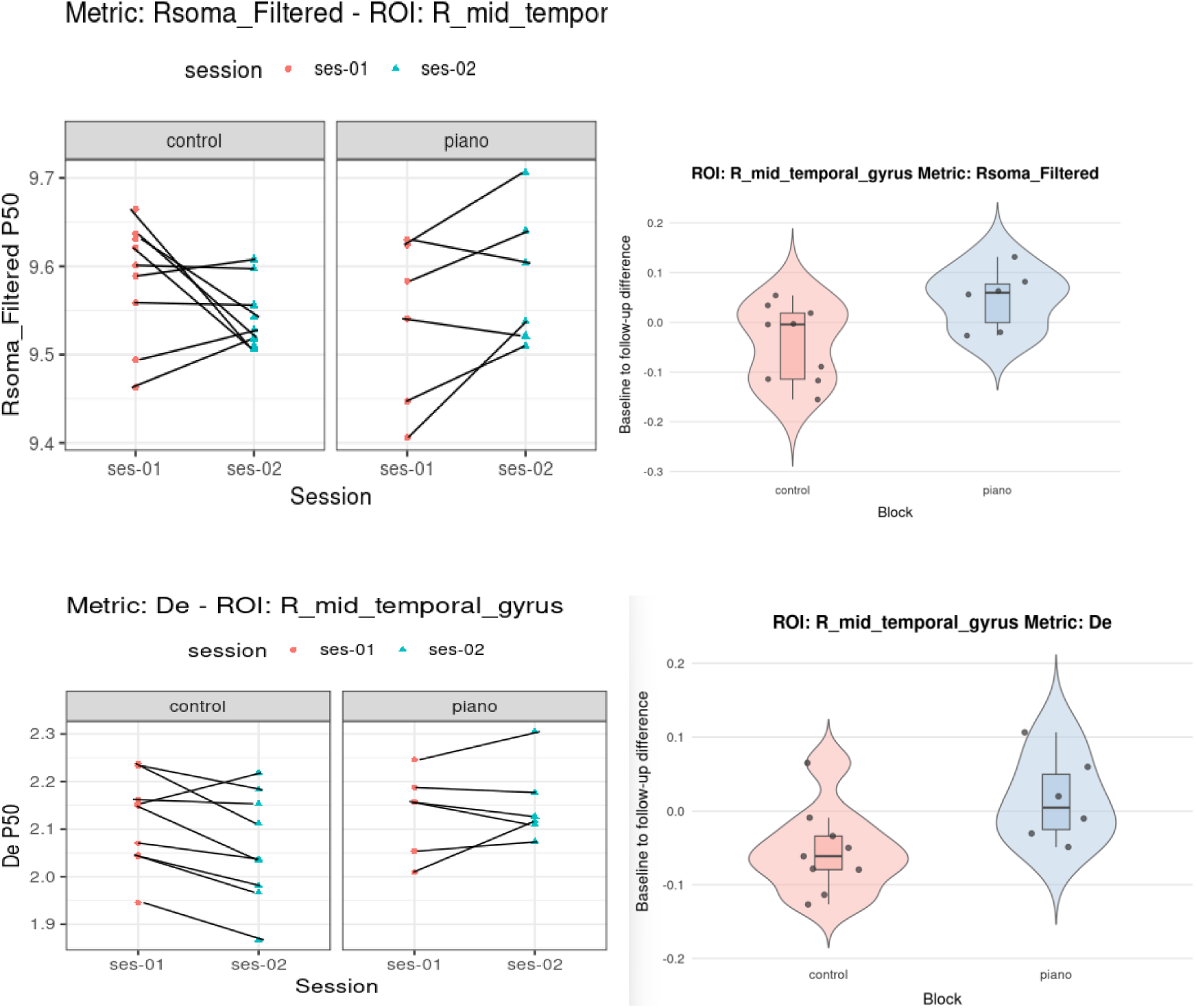
Line and violin plots demonstrating changes in Rsoma and De in right mid temporal gyrus in piano versus control group.

##### Banks of the Superior Temporal Sulcus

An increase for soma size was observed in the right banks of the STS in the intervention group and decrease for the control group, along with a trend for a decrease in white matter microstructure metrics (neurite density measures from SANDI and NODDI), and a decrease in free water metric from NODDI *(Figure 11)*.

**Figure 11:**
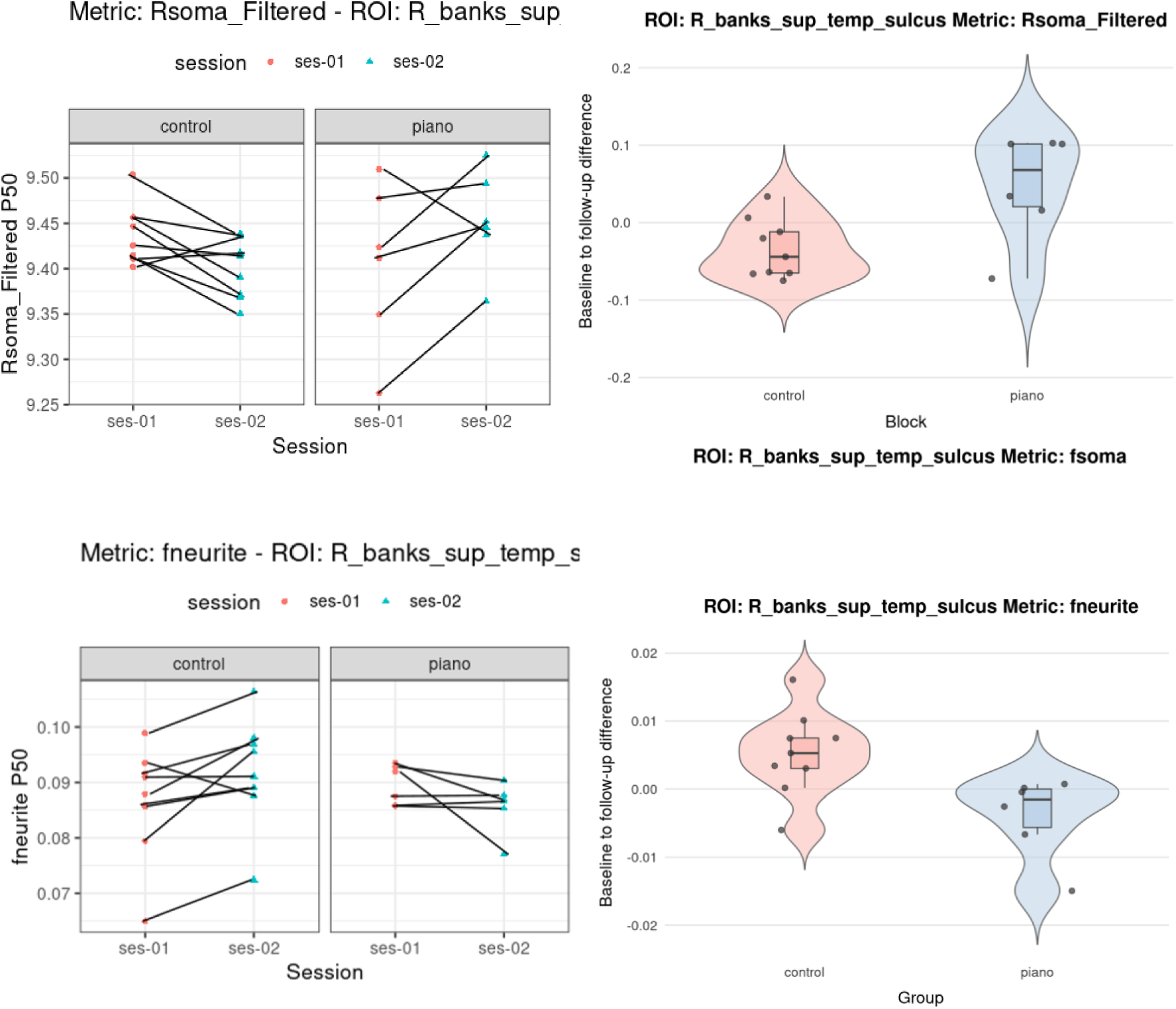
Line and violin plots demonstrating changes in Rsoma and De in right banks of the superior temporal sulcus in piano versus control group.

##### Right Precentral Gyrus

ODI increased for the piano group compared to the control group in the right precentral gyrus, suggesting synaptogenesis in this region associated with motor learning.

##### Summary of grey matter microstructural changes

Overall, exploratory microstructural analyses suggest group differences in changes in neural microstructure in bilateral IFG, bilateral MTG and right banks of STS, and right precentral gyrus. Left IFG and right MTG show changes in soma metrics whilst other regions listed show changes in neurite density or dispersion.

#### 2.3.3 White Matter – Tractography

Two control participants were removed from the tractometry analysis because segments of the CC did not reconstruct. The forest map suggests a trend for greater increase in FA for the piano group *(Figure 13)*.

**Figure 12:**
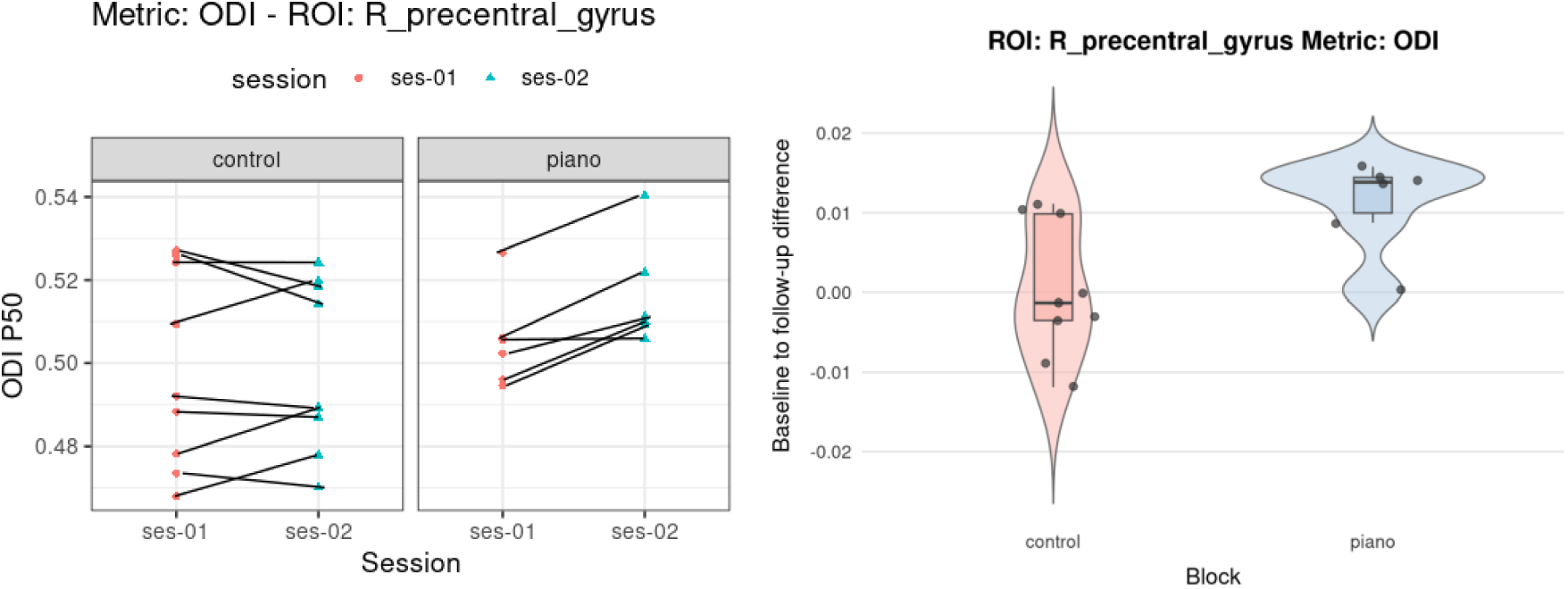
Line and violin plots demonstrating changes in ODI in right precentral gyrus in piano versus control group.

**Figure 13:**
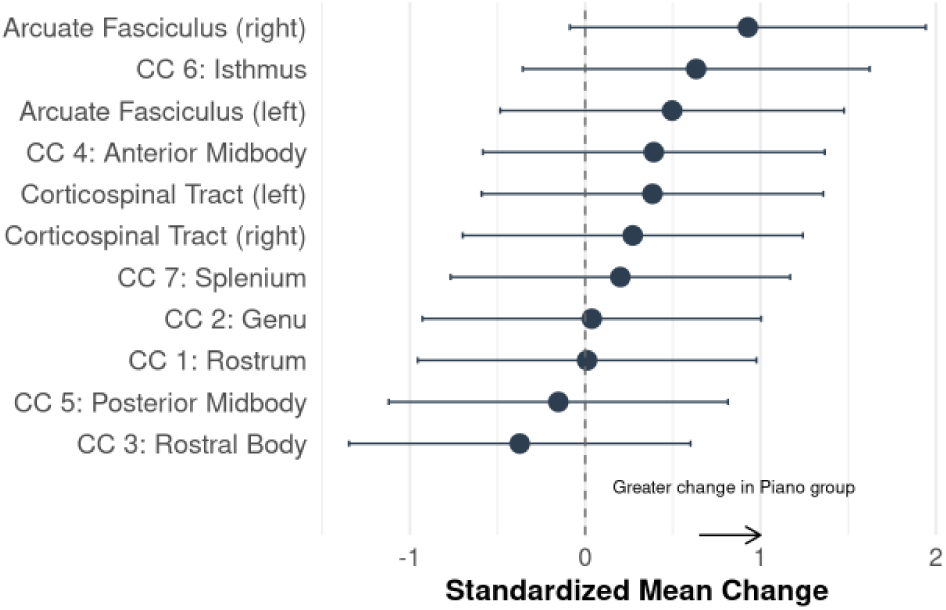
Forest plot demonstrating the differences in fractional anisotropy change between groups from baseline to follow-up across TOIs. CC: corpus callosum.

### 2.4 Unblinding

In the majority of cases (75%), follow-up assessments were carried out by blinded members of the research team. In the cases where unblinded assessor carried out follow-up testing, the reasons were due to illness of a research team member or lack of availablility/ other commitments.

## 3 Discussion

The primary objectives of this study were to assess the acceptability and feasibility of PIANO-Cog, a novel online piano-based cognitive training programme for healthy non-musicians over the age of 50, and to evaluate the feasibility of conducting a fully-powered RCT comparing PIANO-Cog to a passive control group in terms of recruitment, retention and adherence. Secondary objectives were to explore training-induced changes in measures of fluid and executive functions and brain microstructure.

The trial was found to be feasible, achieving a predefined “green” rating for recruitment, retention and adherence measures all scoring >70%. The training was favourably evaluated and participants in the piano group reported practicing an average of 30 hours over the course of the 8-week intervention, exceeding the requested 20 hours. These findings support the acceptability of remote piano training as a cognitive intervention for older adults.

### 3.1 Cognitive Outcomes

Preliminary observations suggest that participants in the piano group improved performance on letter verbal fluency compared to the control group and showed a trend for improved category fluency in line with (72). Trends were also observed for inhibitory control (measured using the Go/No-Go task) consistent with findings from our meta-analysis (18) and working memory (measured using the N-Back task), in line with (73) and (33). In contrast with previous piano training studies in ageing, no changes in processing speed measured using Symbol-Digit Test (36,72–74), or in attention switching using TMT-B were observed (36,75,76). However, prior studies reporting benefits in these domains had larger sample sizes and employed longer interventions (at least 16 weeks), with the exception of (72), which was an intensive course of 3 hours per day over 2 weeks.

The positive change in letter fluency suggests that short-term musical training may target executive and language-related processes beyond the effects of generalized improvements in processing speed as previously suggested (72). This is promising considering that verbal fluency is thought to rely on executive processing and memory (1) mediated by frontal and temporal cortex which are known to be disproportionately affected by ageing (2). However further research in a fully-powered RCT is needed to replicate these findings and establish whether cognitive benefits are correlated with brain microstructural changes.

### 3.2 Microstructure

Preliminary observations suggested microstructural changes in the IFG. In the left IFG (Broca’s area), the piano group showed an increase in *fsoma* alongside a decrease in *fextra* compared to the control group, suggesting a change in the proportion of the estimated MRI signal to be coming from soma, potentially related to glial swelling. In the right IFG, piano participants showed a slight increase or maintenance of *fneurite* (SANDI) and neurite density index (NDI; NODDI), whereas the control group tended to show a decrease over time, suggesting preservation of neurite-related metrics in the piano compared to the control group, which tend to decline in ageing (77).

Preliminary changes were also observed in the bilateral mid-temporal gyri. In the left MTG, ODI decreased in the control group but remained stable or increased in the piano group, alongside a reduction in fractional anisotropy (FA), potentially reflecting more complex dendritic fanning that may lead to reduction in FA. In the right MTG and right banks of the STS, *Rsoma* increased in the piano group and decreased in the control group, again suggesting possible glial swelling in this region. Right MTG also showed a decrease in extracellular diffusivity (De) in controls but remained stable in piano participants. Together, these patterns suggest group differences in temporal lobe microstructural metrics over time, consistent with previous reports of volumetric increases in the temporal lobe following 6-months of piano training (32).

Finally, there were preliminary observations of piano-related increases in ODI in the right precentral gyrus (motor cortex), potentially due to greater dendritic fanning associated with increased use of the left hand in predominantly right-handed participants.

At the whole-brain level, piano training related reductions in MD of right-hemisphere white matter were observed, consistent with above predominantly right-lateralised microstructural changes. This may suggest training-related adaptations in right-lateralized white matter microstructure following increased non-dominant left-hand activity (ref).

It should be noted that these microstructural changes were observed following a relatively short 8-week intervention with a limited sample size. Replication is needed in a fully-powered RCT. Future studies should also take temporal dynamics into account as preliminary research suggests that training-induced changes in soma metrics return to baseline within 24 hours, possibly reflecting homeostatic processes, whereas neurite measures may be more stable over time and reflect synaptic remodelling (78).

### 3.3 Limitations and Future Directions

As this was a feasibility study, it was not fully-powered and only 59% of participants were eligible for MRI scanning on the Connectom scanner. Nevertheless, the data showed clear signatures in behavioural and microstructural measurements providing preliminary evidence that music-based approaches may be helpful for slowing neural and cognitive decline in normal ageing.

Although stratified allocation was used to balance age and sex across groups, and no baseline differences were observed in global cognitive ability, years of education, music perception scores on microPROMS or any cognitive measure, the piano group scored higher on the TOPF, suggesting higher estimated verbal intelligence at baseline. This may have influenced how piano participants engaged with the training material and applied their own learning strategies, and should be more carefully controlled in future studies.

Another potential limitation of this format of self-guided intervention is that recruitment may be biased towards individuals who are motivated by self-guided musical training. Additionally, successful engagement with the programme likely requires the ability to follow instructional videos and a level of self-discipline to for structured practice using a metronome.

Importantly, we did not observe any overall changes in musical perception skills measured using the microPROMS (45), which may suggest that although remote training could improve cognition due to executive demands, improvements in musical perceptual abilities may require longer or more individualised in-person instruction.

### 3.4 Conclusion

PIANO-Cog was found to be an acceptable remote cognitive intervention for healthy ageing and feasibility measures of recruitment, retention and adherence suggest a fully-powered RCT is feasible. In line with previous work, we report preliminary observation of improvements in verbal fluency after 8-week piano intervention and changes in brain microstructure in brain regions associated with auditory, motor and executive processes highlighting the potential of advanced diffusion MRI to identify training-induced plasticity mechanisms.

## Funding

This work was supported by an Open PhD studentship from the School of Psychology Cardiff University and by a National Institute for Health Research (NIHR) and Health and Care Research Wales (HCRW) Advanced Fellowship to CM-B (grant number: NIHR-FS(A)-2022).

## Competing Interests

The authors declare that they have no competing interests.

## Acknowledgements

We would like to sincerely thank Dr. Marco Palombo for his expertise with SANDI model interpretation, our research assistants, Beate Galoburda and Katrin Powell, for their help with data collection, as well as all our participants who gave their time to testing, scanning and learning piano.

## Notes

### Competing Interest Statement

The authors have declared no competing interest.

