## supplementary material for "Feasibility of PIANO-Cog for older adults: A randomised controlled pilot trial exploring changes in cognition and brain microstructure"

### 1.1 Age Distribution of Sample

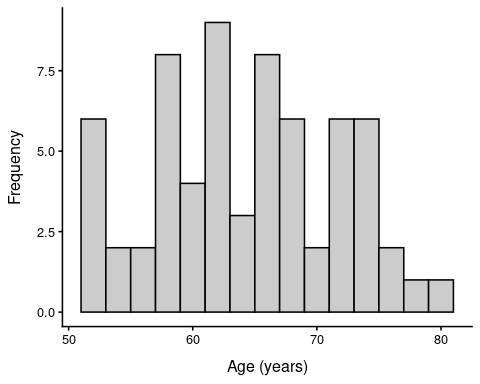

### 1.2 Tests for Group Differences at Baseline

| Test | W | p | Control Median | Piano Median |
| --- | --- | --- | --- | --- |
| Digit_Symbol | 221.5 | 0.671 | 52.00 | 52.50 |
| Digit_span_Backward | 177.0 | 0.194 | 7.00 | 7.50 |
| Digit_span_Forward | 144.0 | 0.023 | 11.00 | 13.00 |
| LDCR | 252.5 | 0.775 | 12.00 | 13.00 |
| LDFR | 249.5 | 0.831 | 12.00 | 12.00 |
| SDCR | 258.0 | 0.676 | 13.00 | 12.50 |
| SDFR | 236.5 | 0.943 | 12.00 | 12.00 |
| TICS | 147.0 | 0.105 | 29.50 | 31.00 |
| TMTA | 285.0 | 0.294 | 27.11 | 25.24 |
| TMTB | 290.0 | 0.243 | 59.34 | 48.52 |
| TOPF | 79.0 | 0.001 | 57.50 | 65.00 |
| Total_CVLT | 186.0 | 0.206 | 51.50 | 56.00 |
| Verbal_Fluency_CF | 142.0 | 0.147 | 44.00 | 47.00 |
| Verbal_Fluency_CS | 183.0 | 0.180 | 15.00 | 16.50 |
| Verbal_Fluency_LF | 182.5 | 0.732 | 42.00 | 50.00 |
| microPROMS | 211.0 | 0.830 | 8.75 | 9.00 |
| Note. Wilcoxon rank-sum tests comparing baseline scores between Control and Piano groups. | | | | |

## 1.3 Go/No-Go

#### 1.3.1 Hits

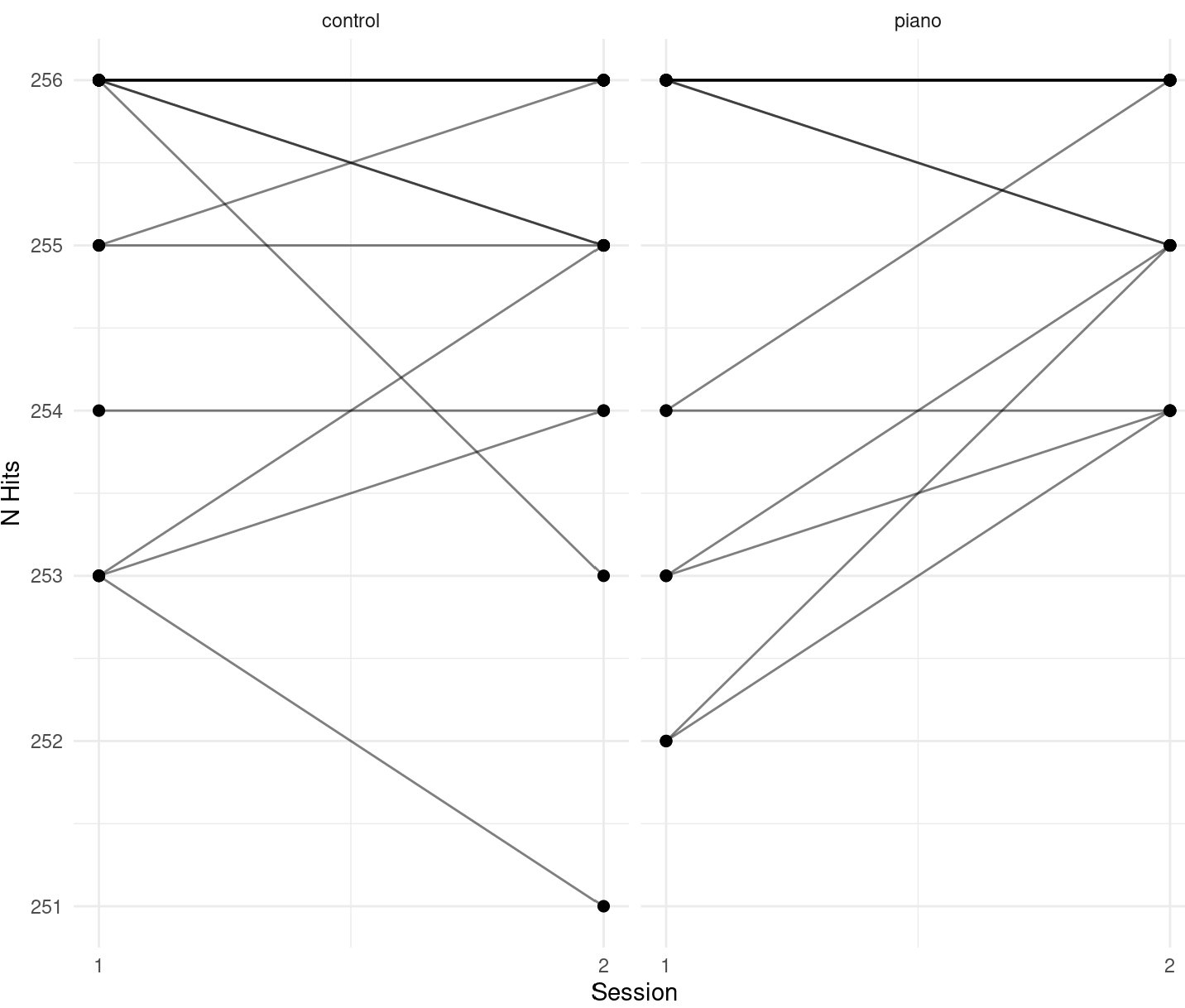

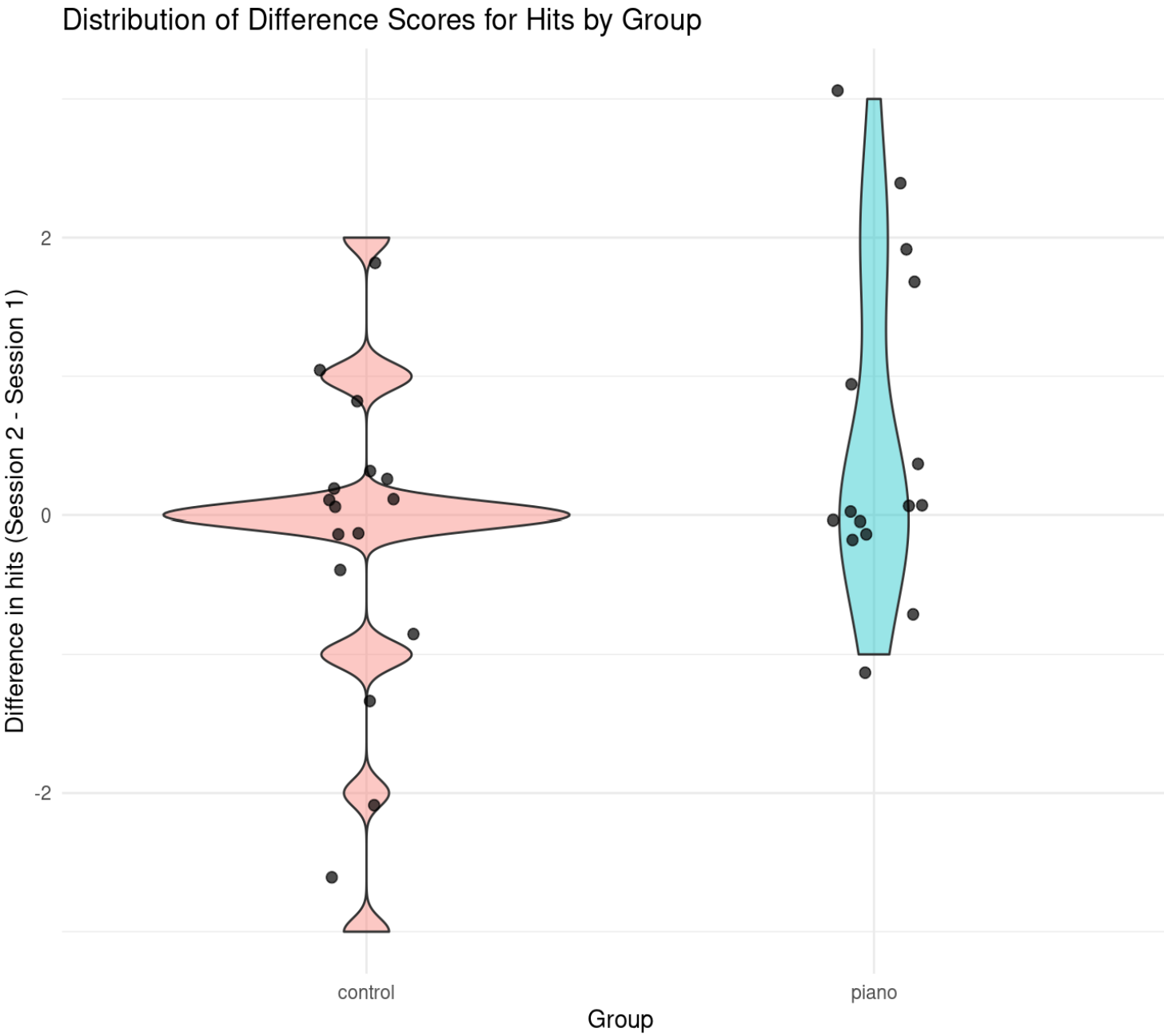

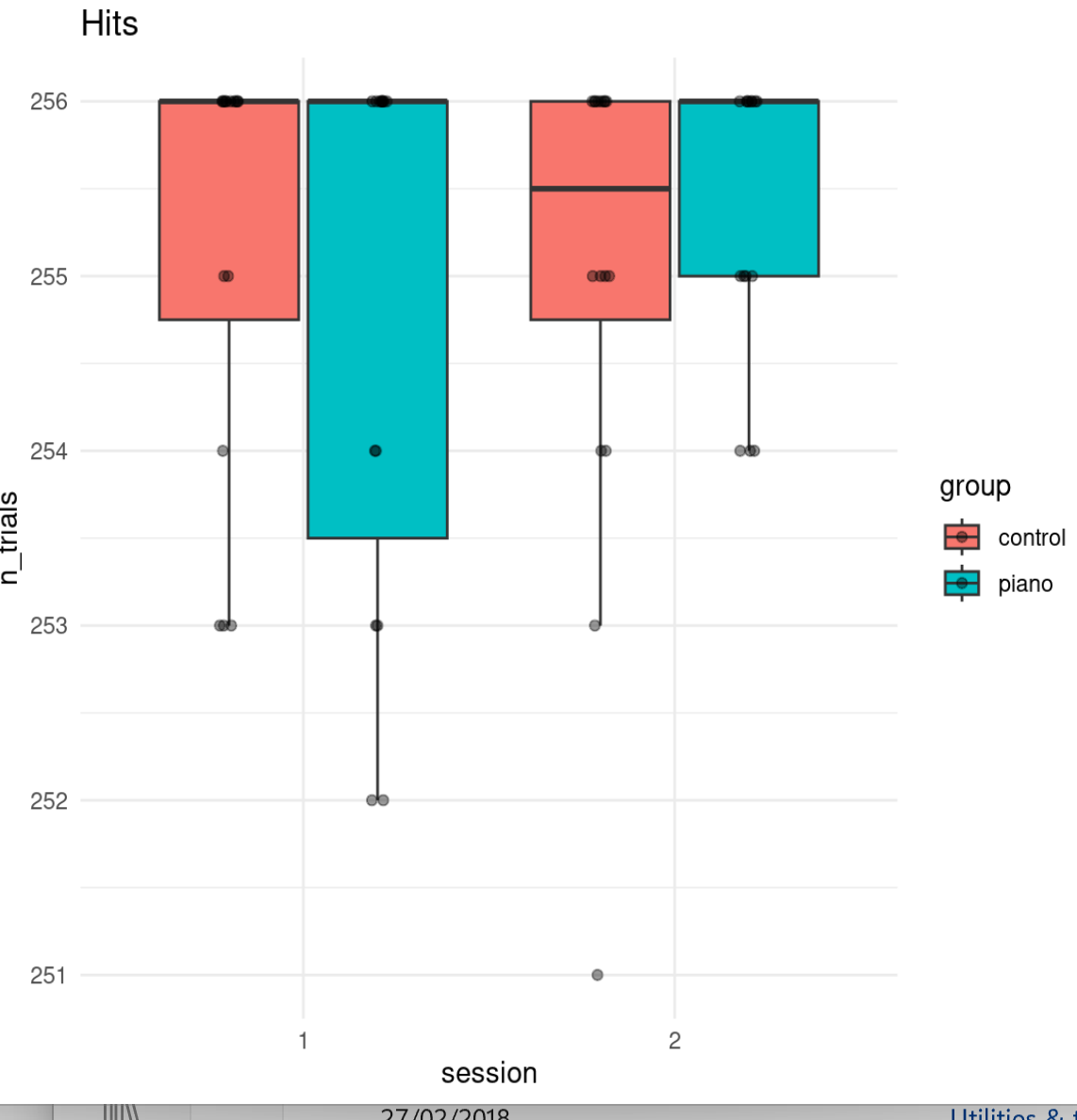

#### 1.3.2 Misses

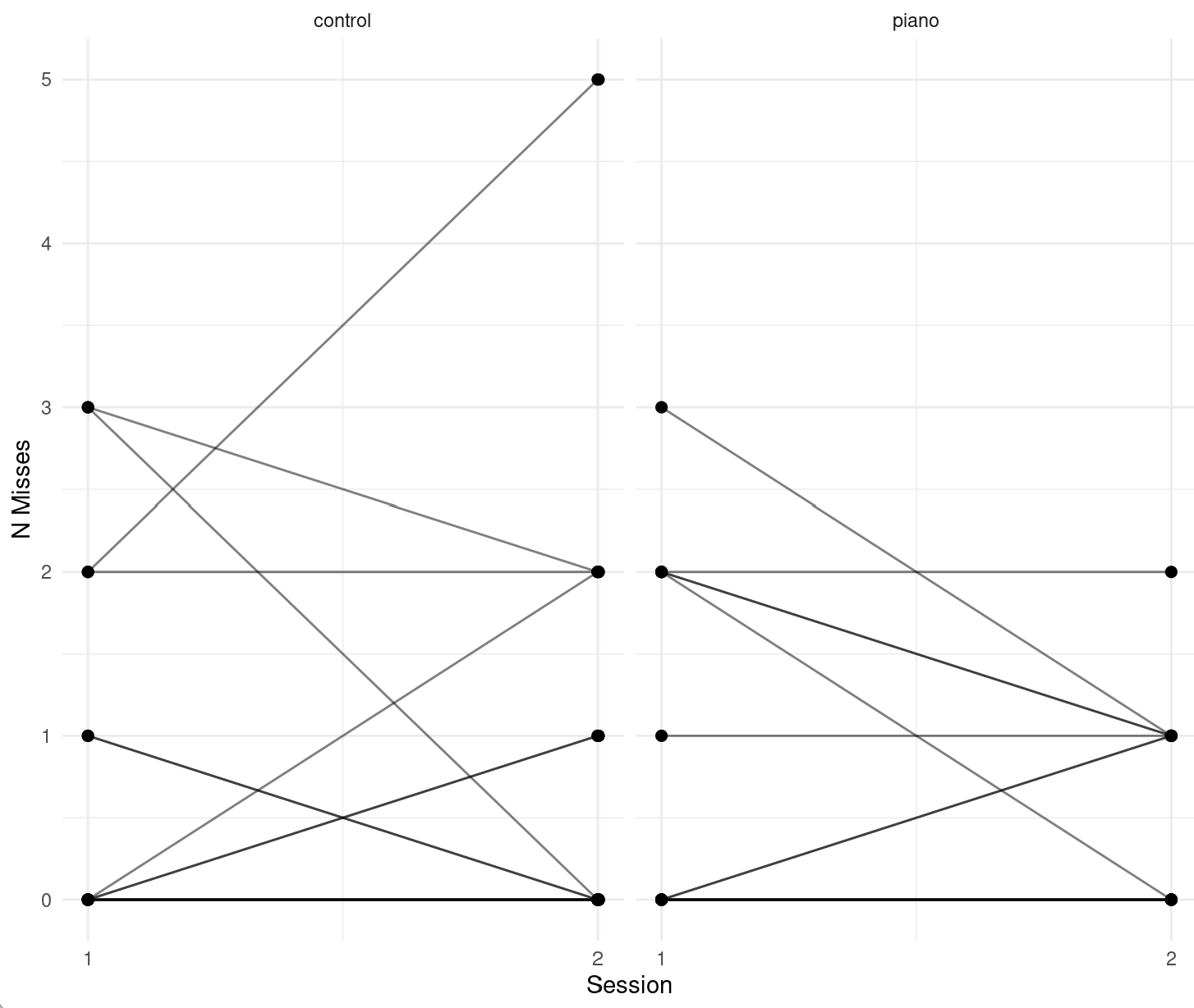

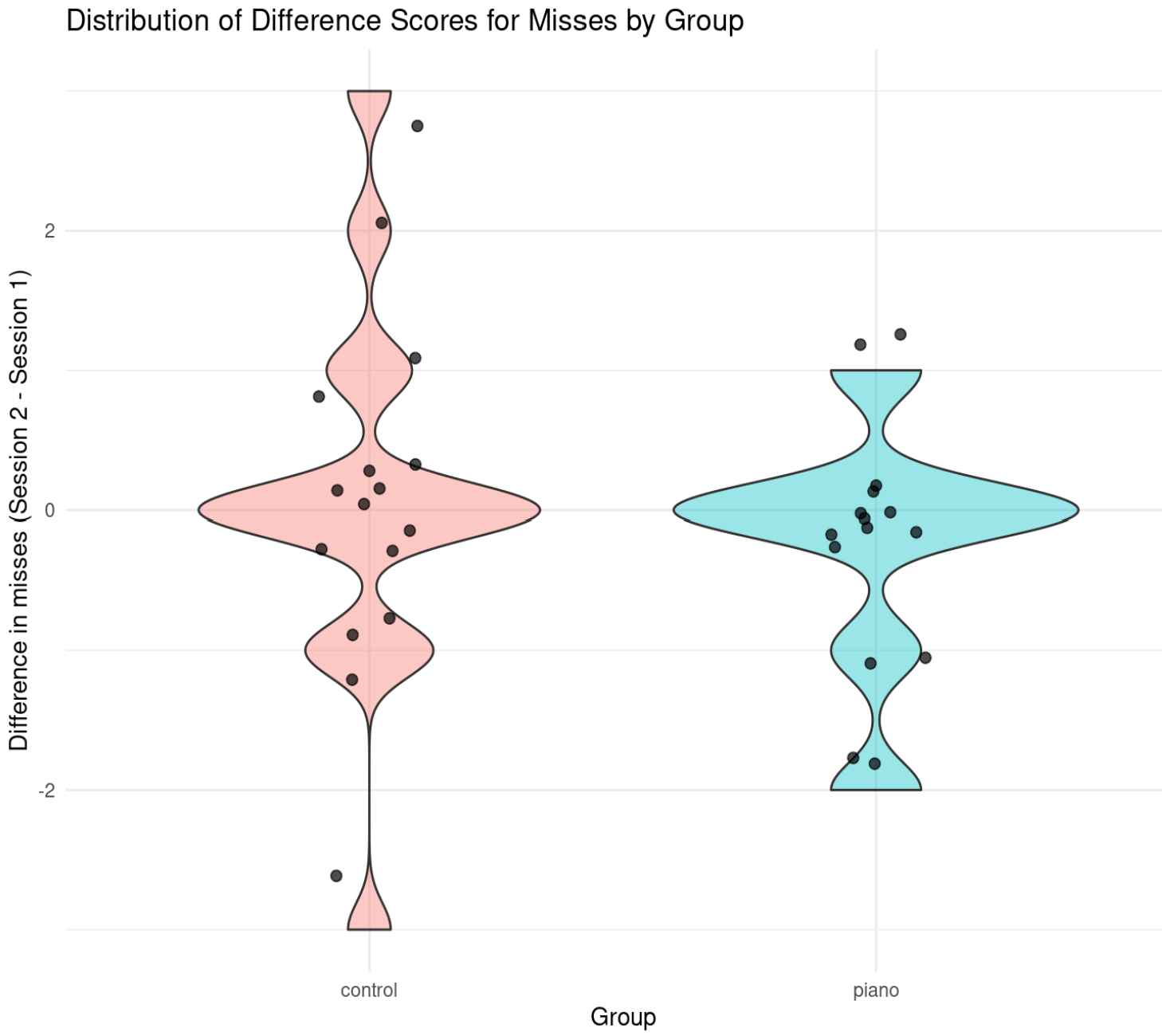

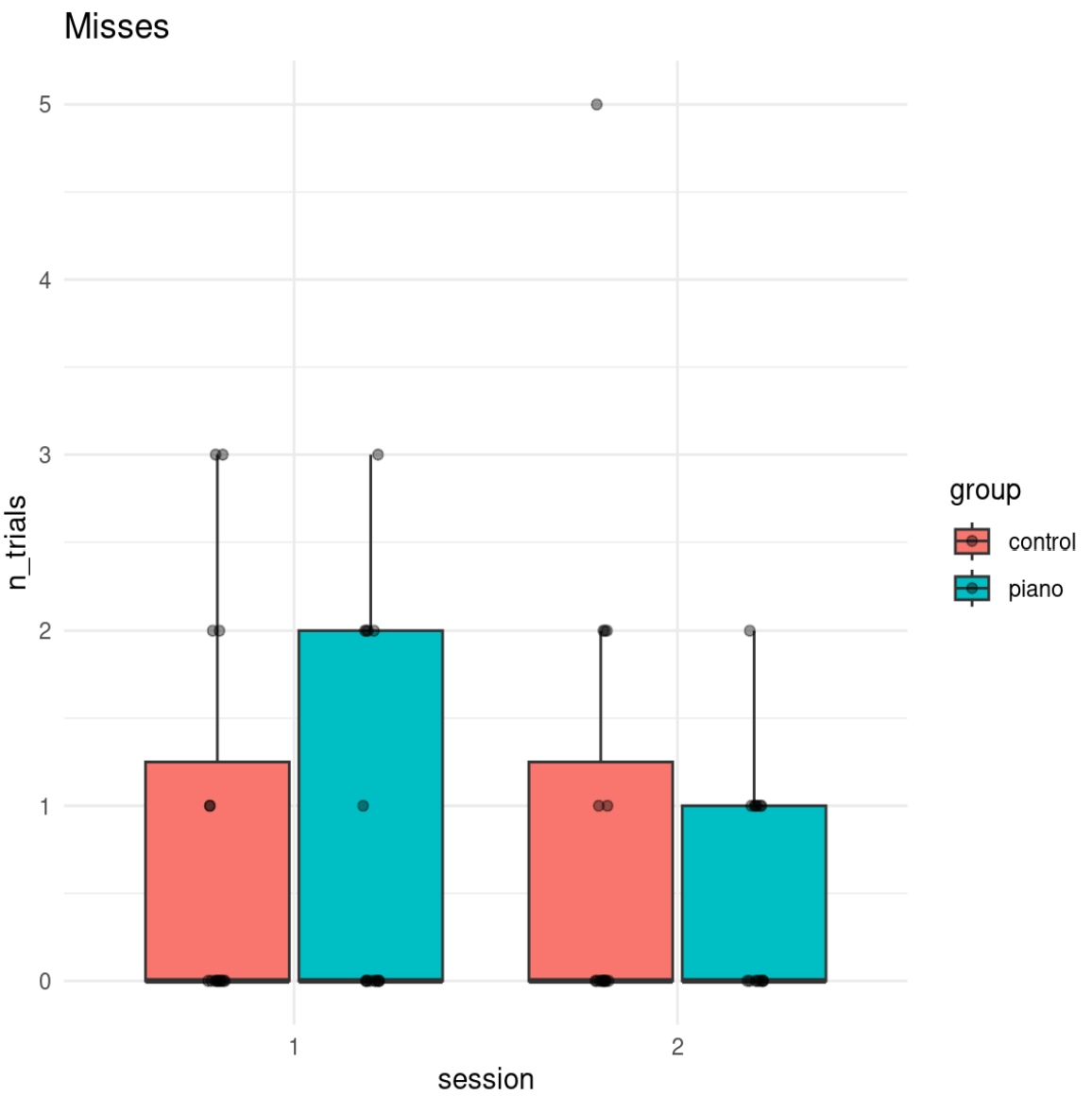

#### 1.3.3 False Alarms

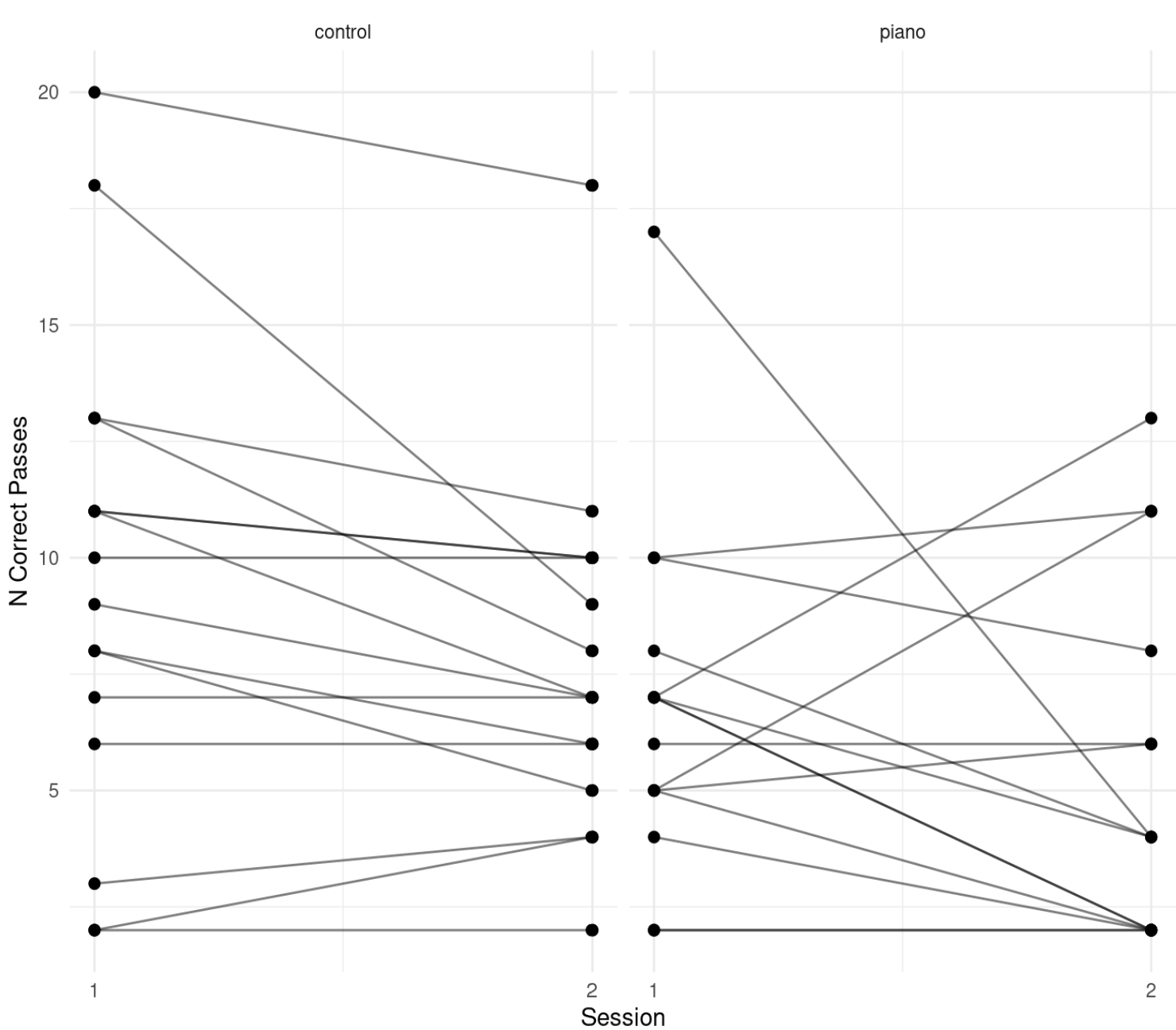

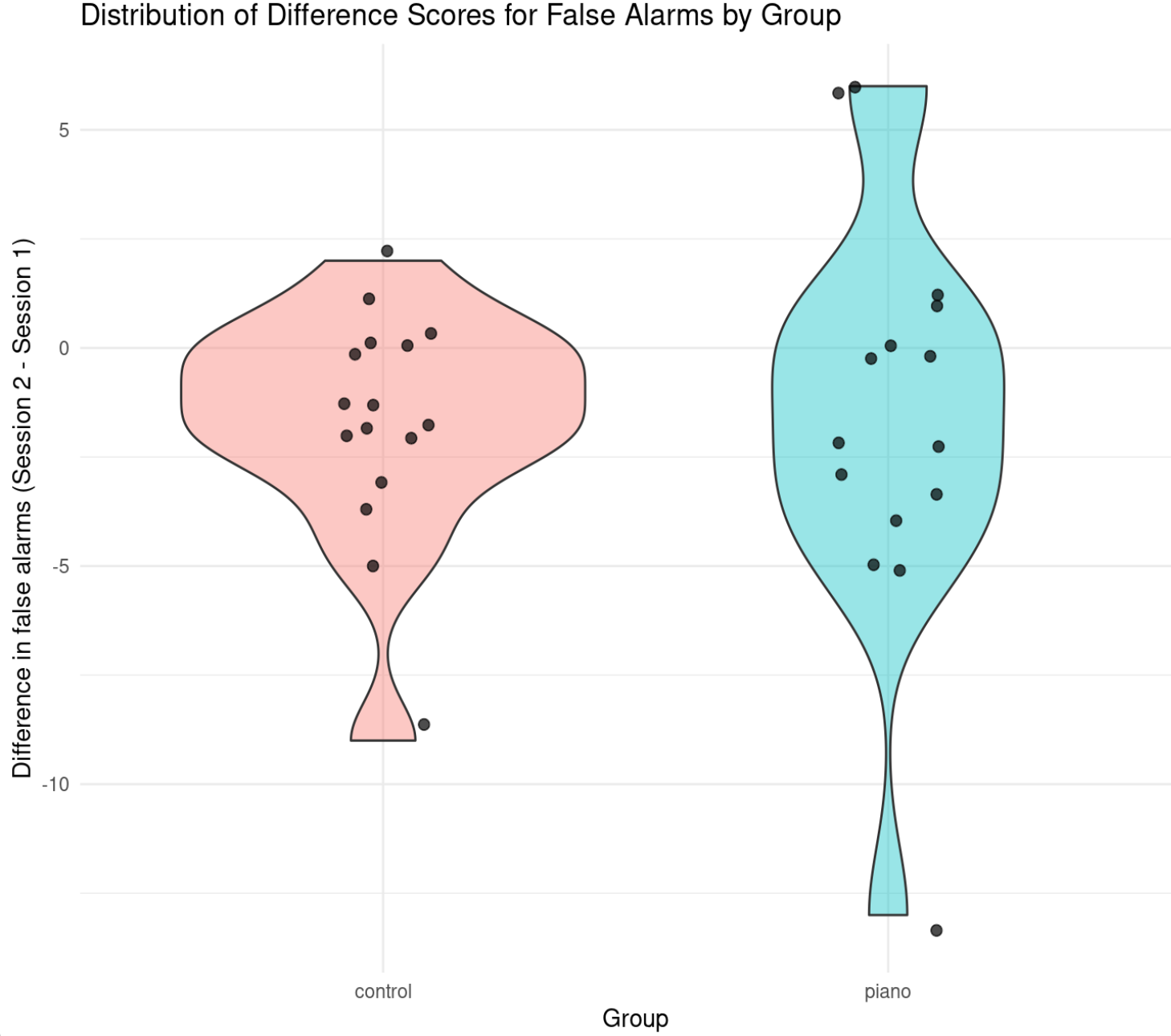

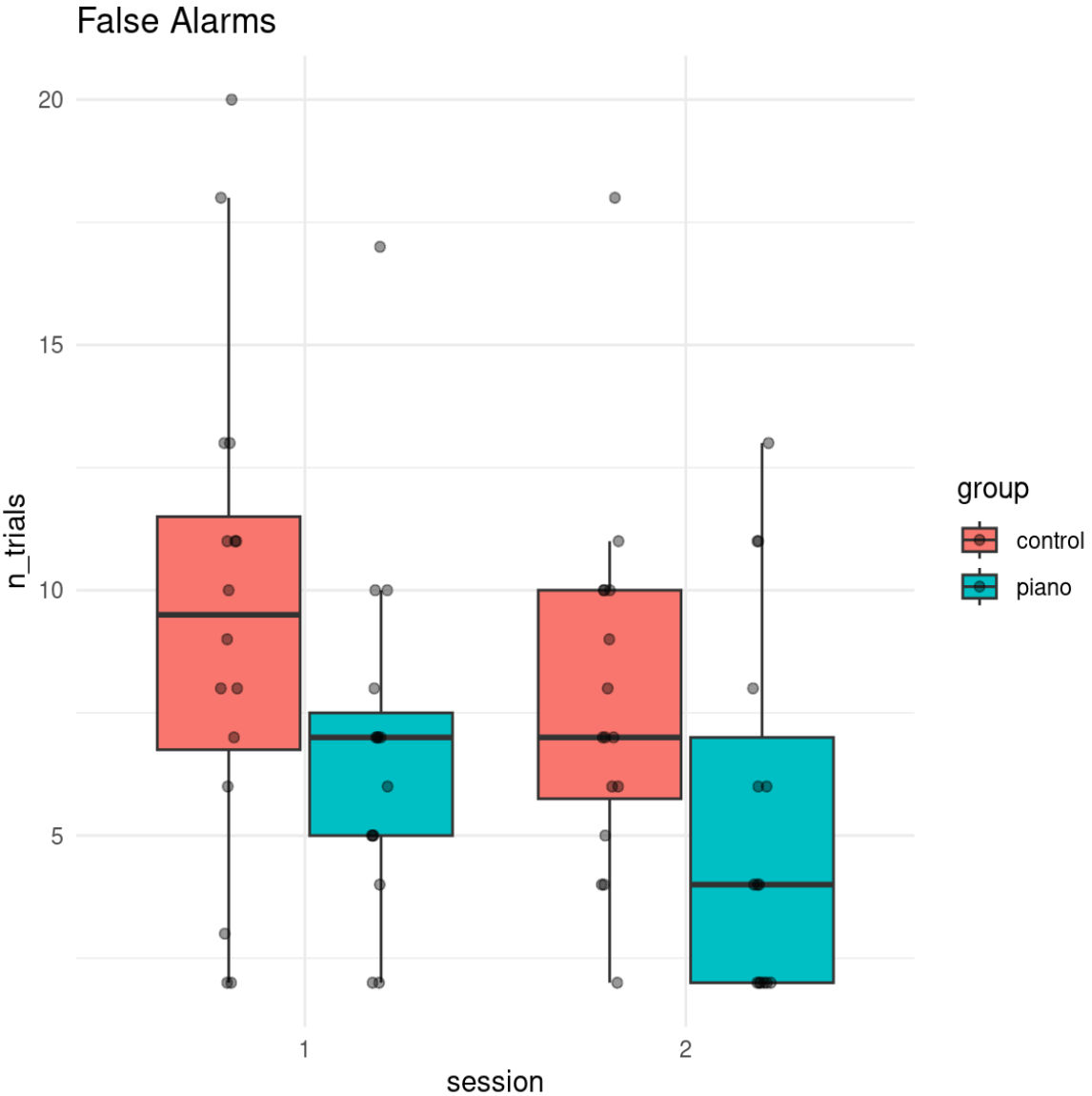

#### 1.3.4 Correct Passes

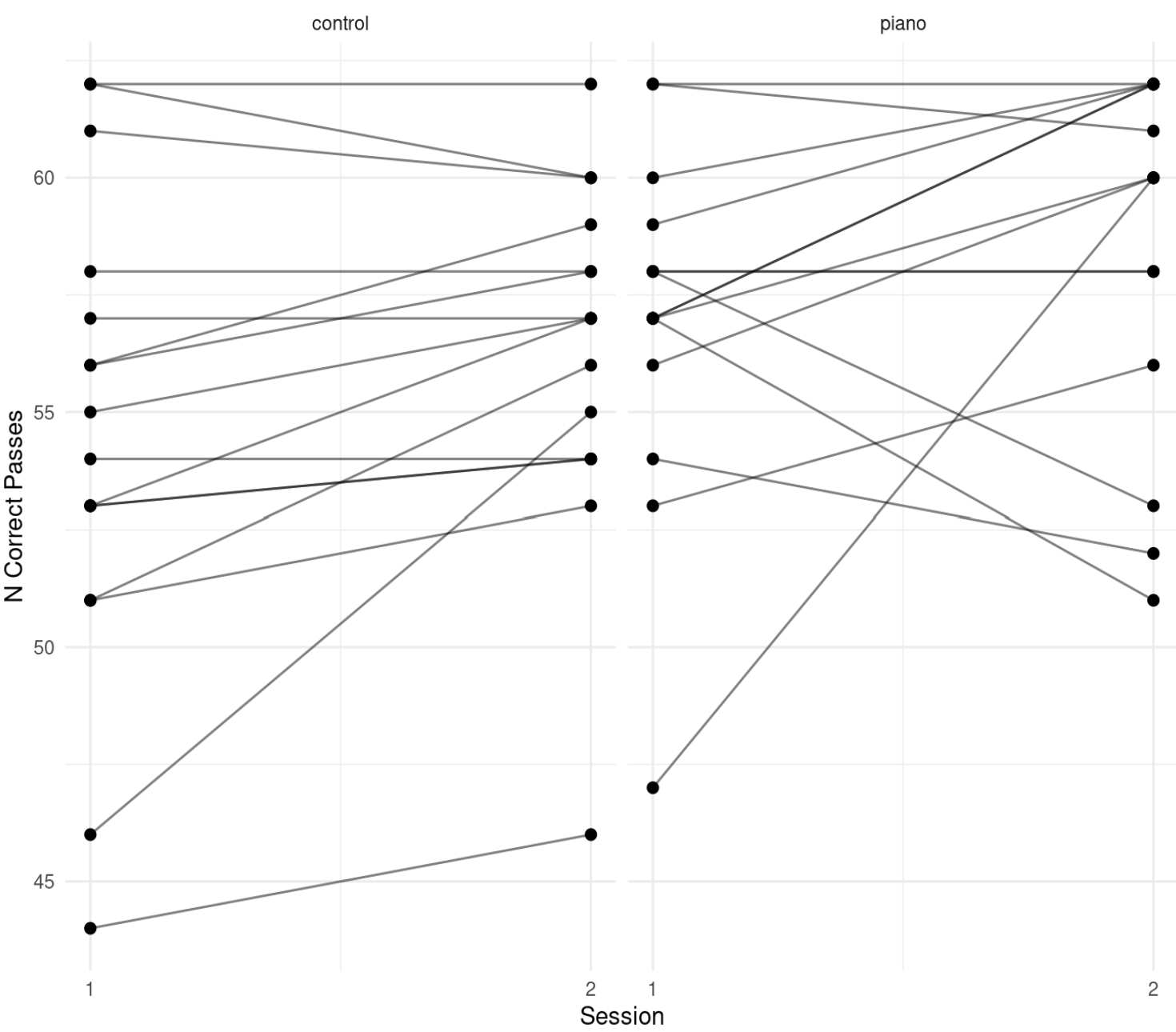

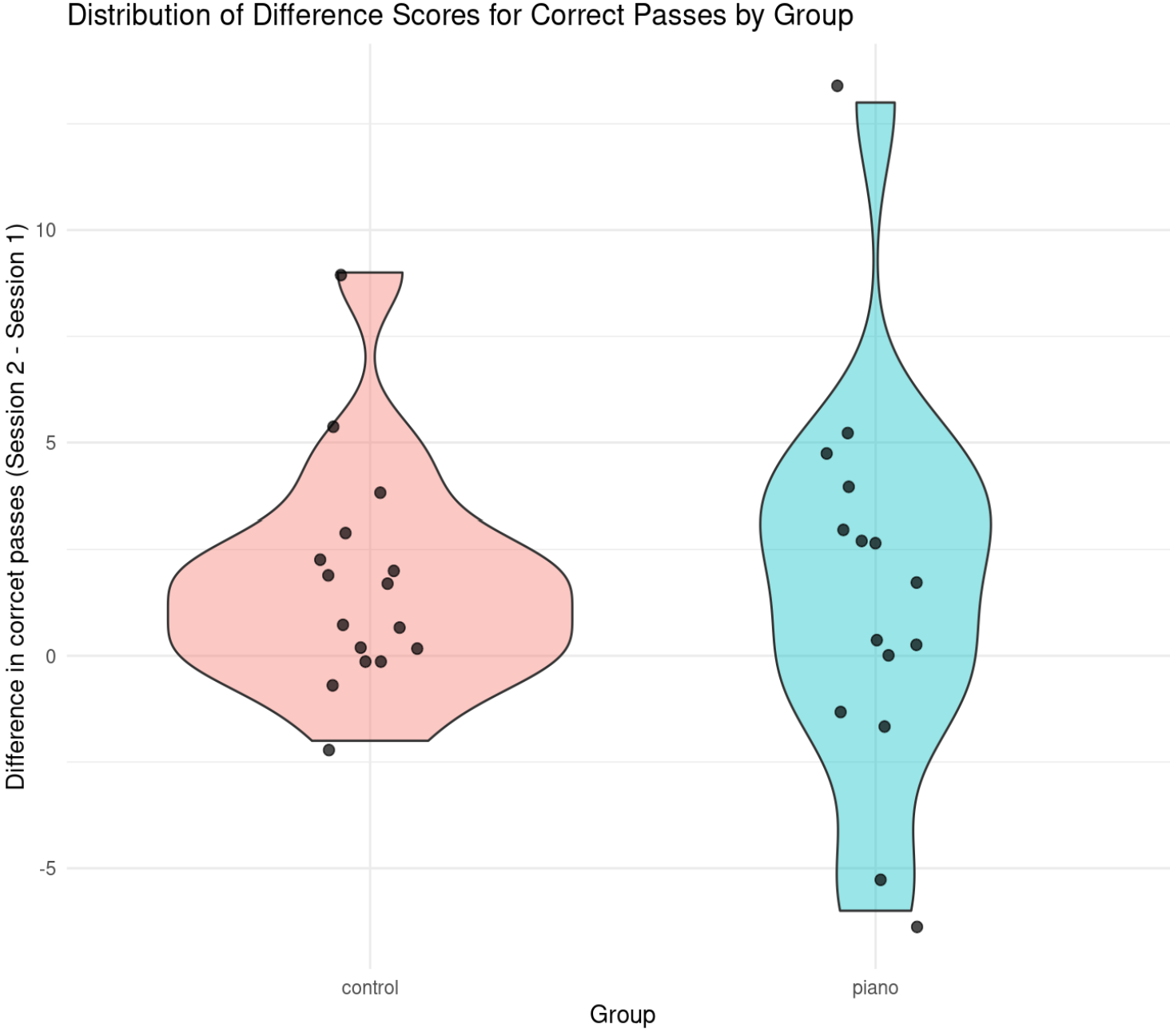

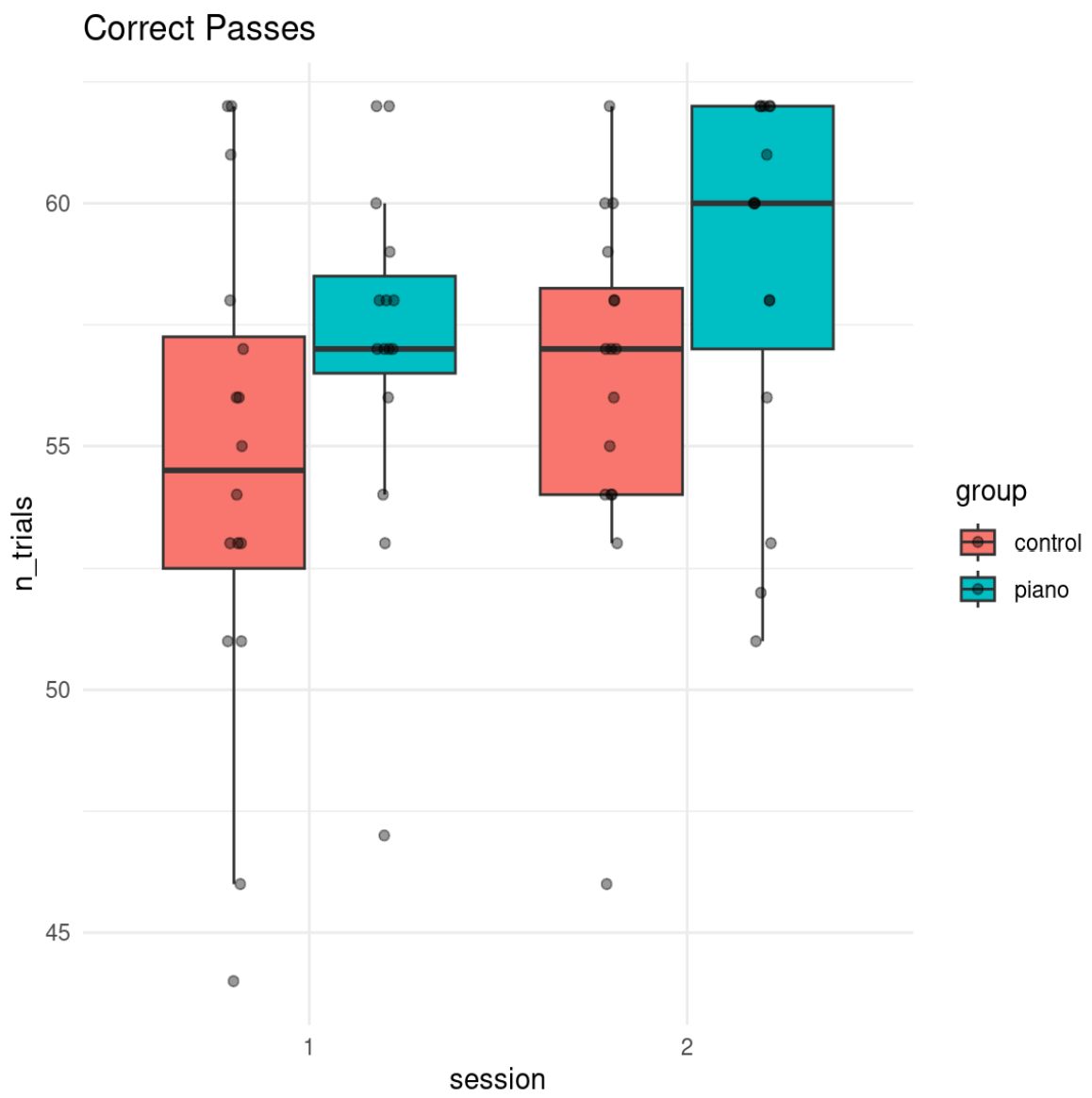

### 1.4 N-back

#### 1.4.1 Dual-Task 2-Back

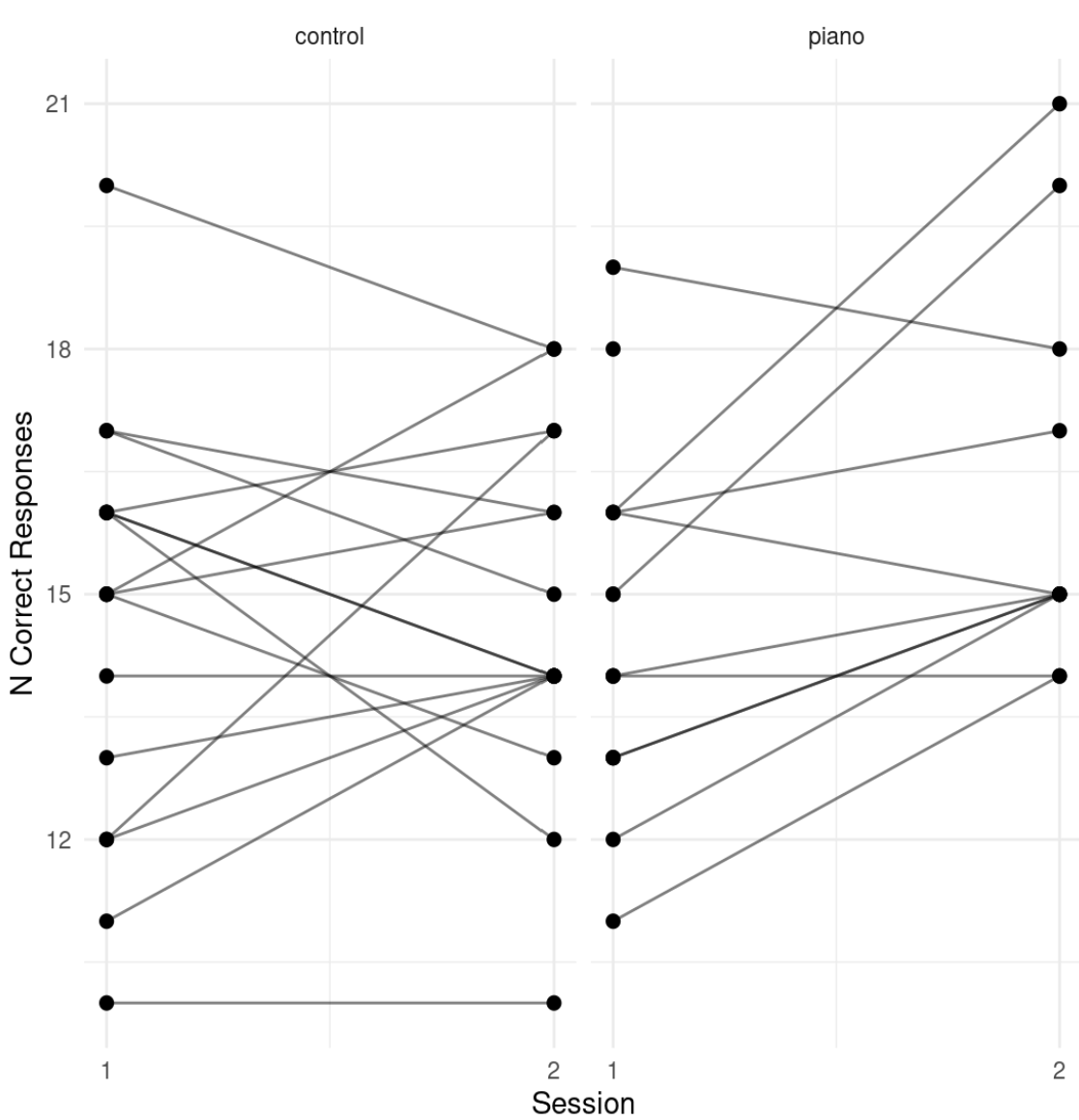

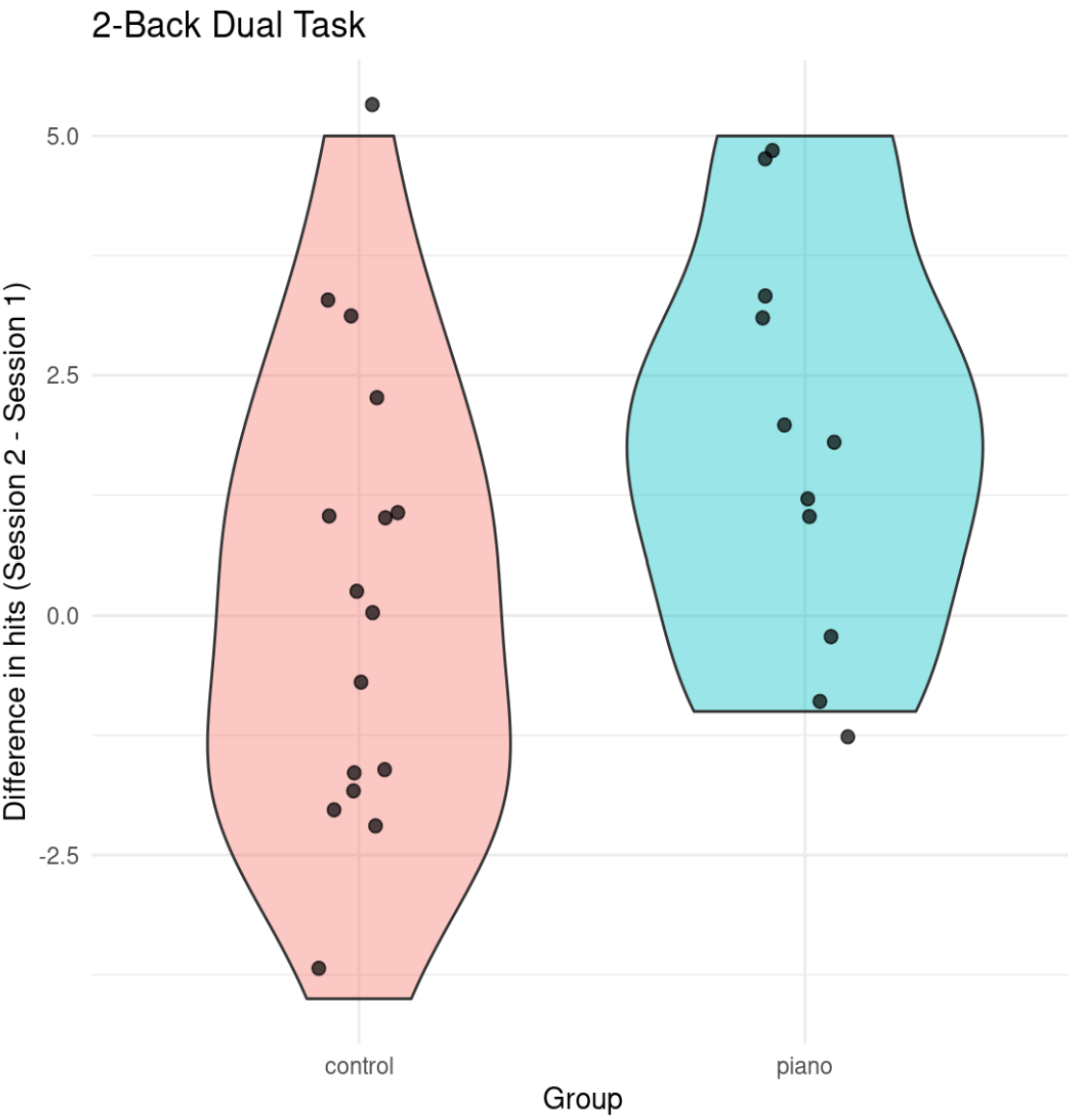

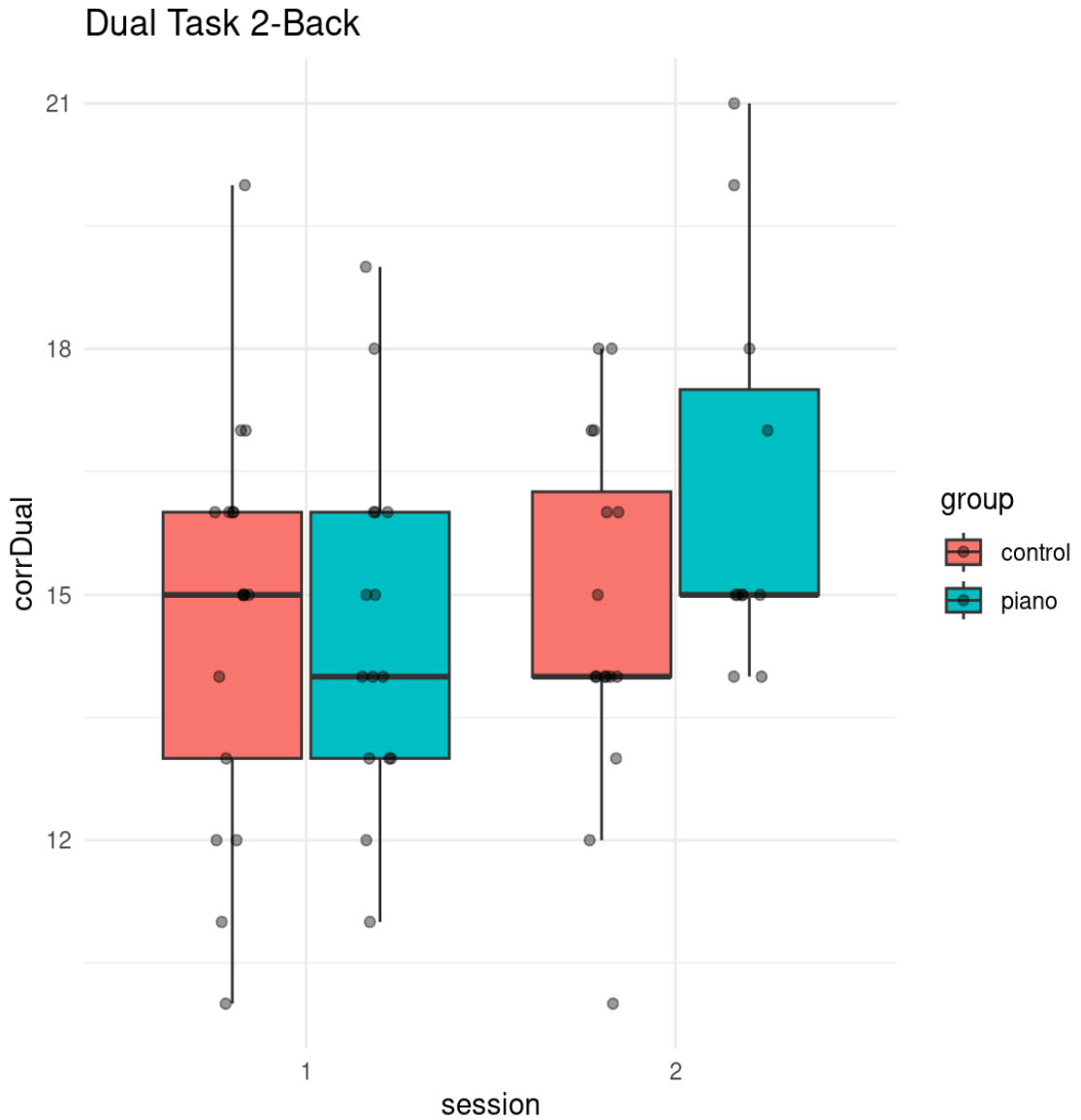

#### 1.4.2 Letter 2-Back

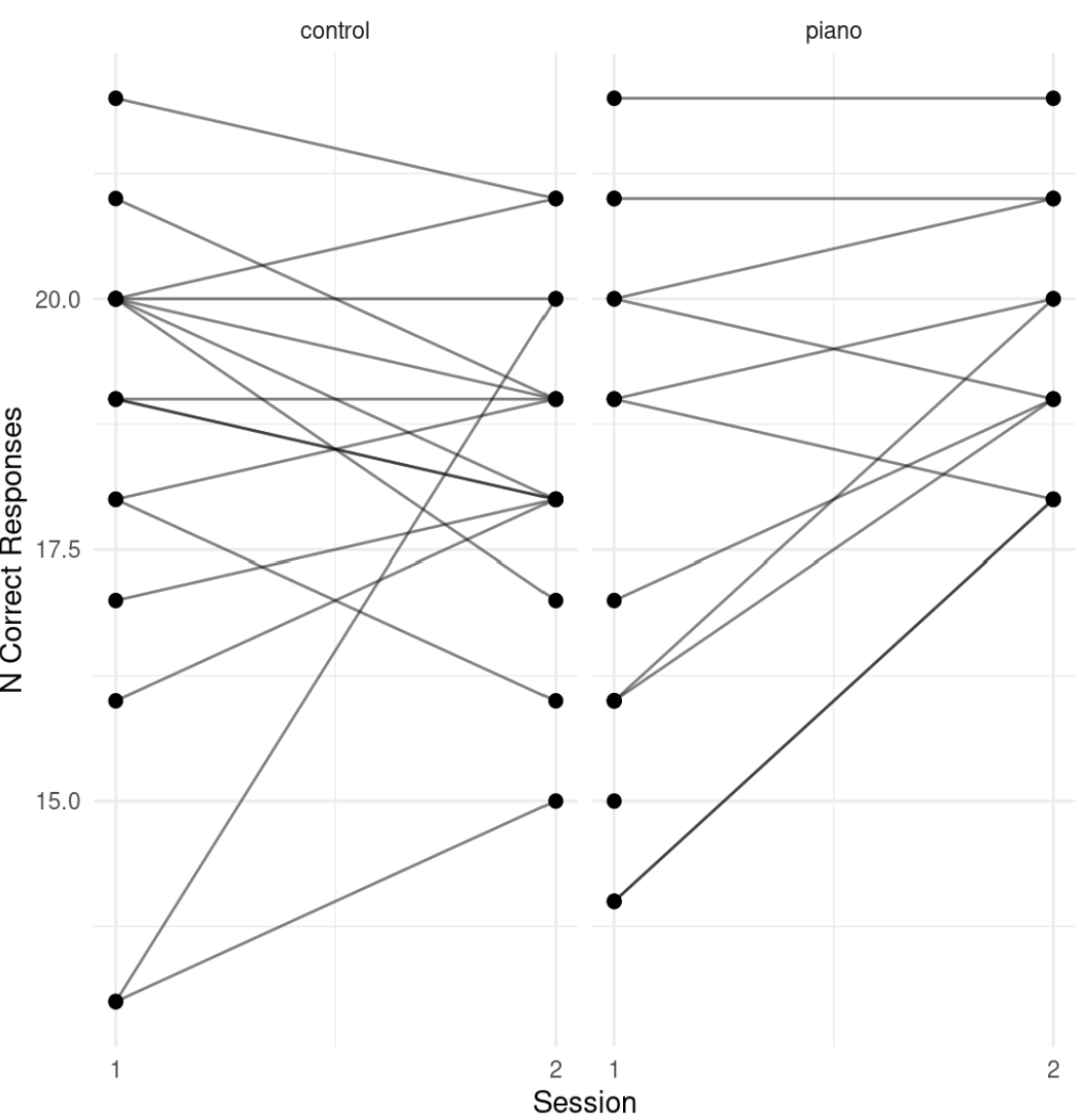

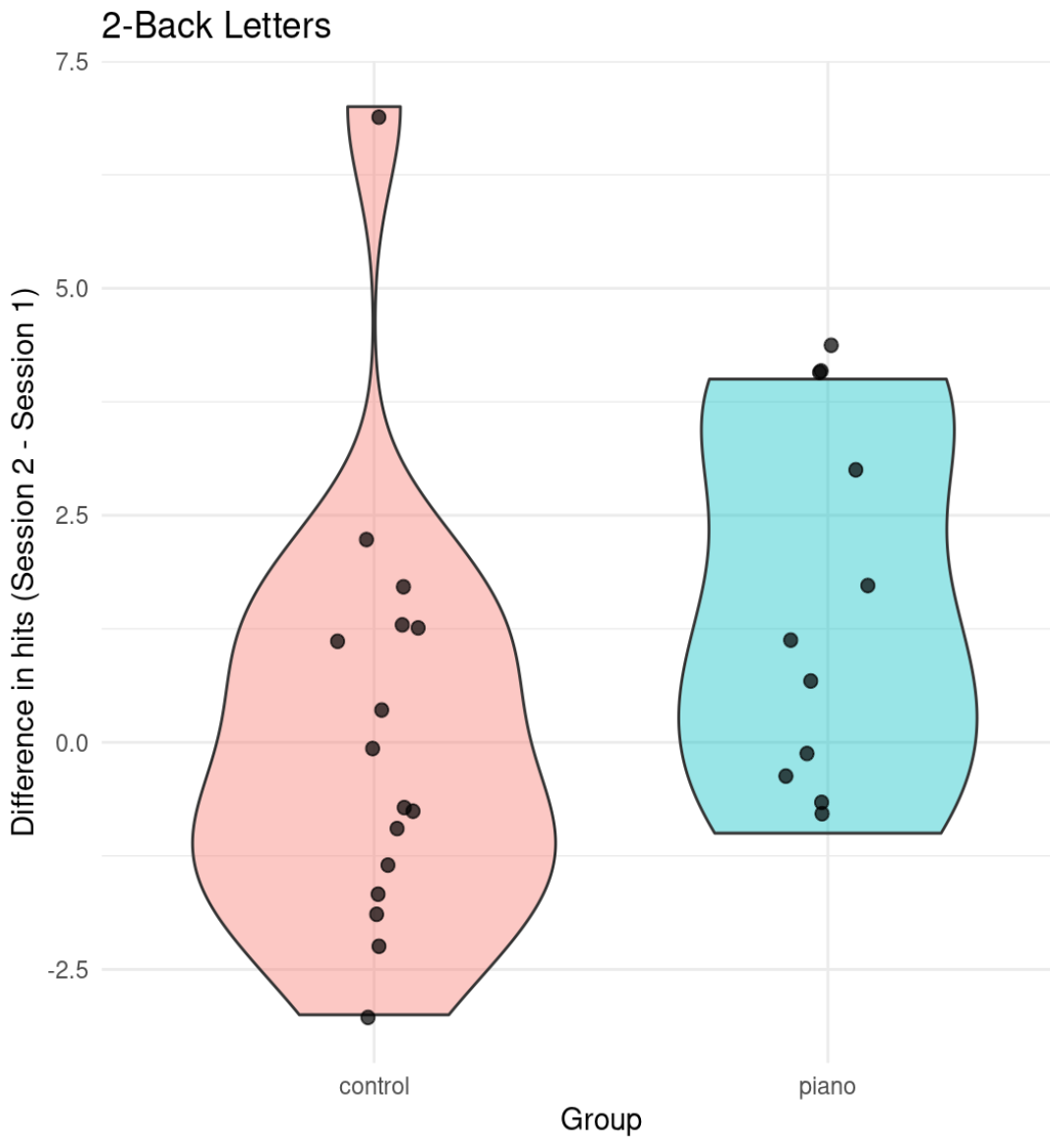

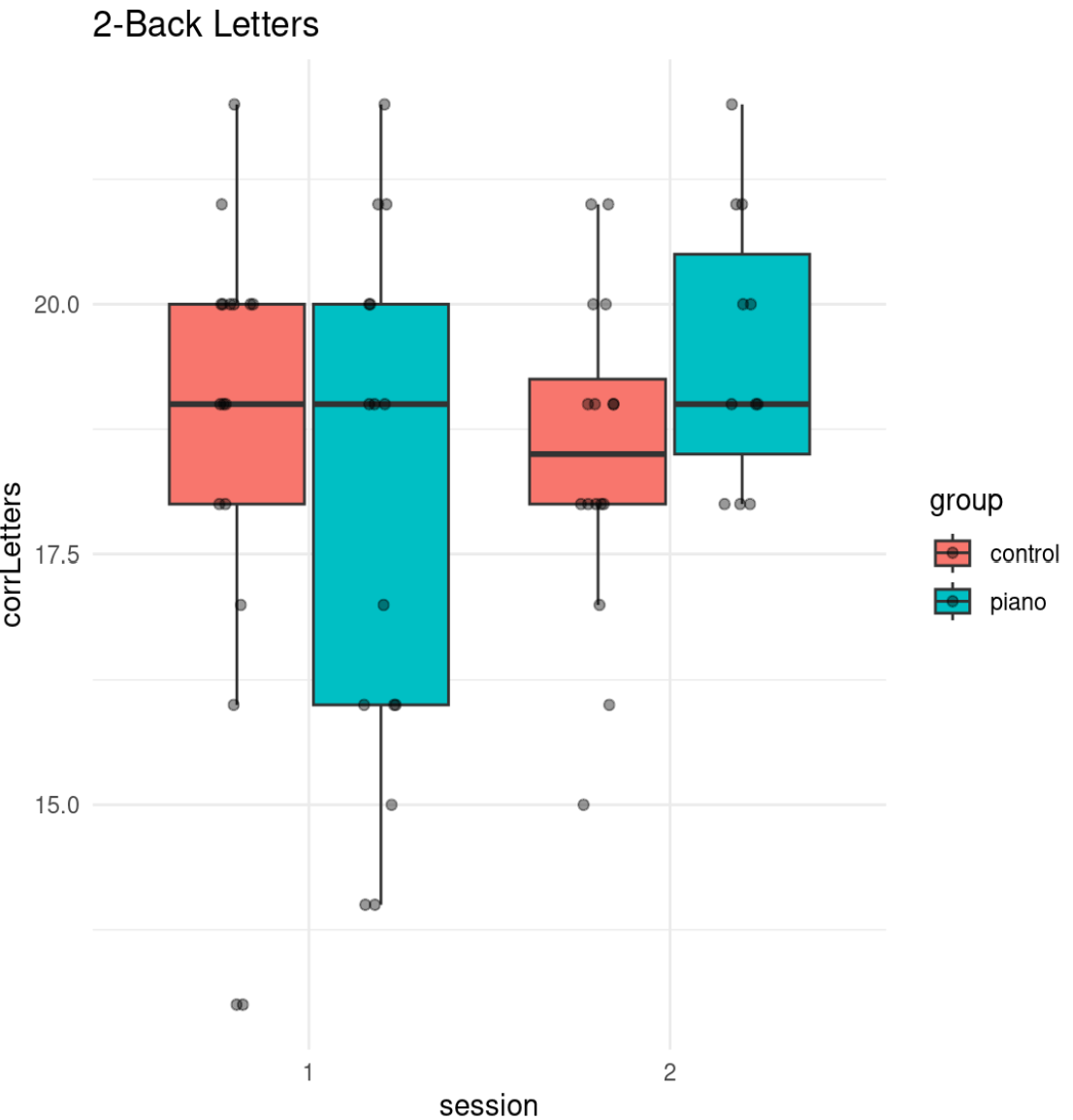
